# Agent-based vs. equation-based multi-scale modeling for macrophage polarization

**DOI:** 10.1101/2022.06.20.496801

**Authors:** Sarah B. Minucci, Rebecca L. Heise, Angela M. Reynolds

## Abstract

Macrophages show high plasticity and result in heterogenic subpopulations or polarized states identified by specific cellular markers. These immune cells are typically characterized as pro-inflammatory, or classically activated M1, and anti-inflammatory, or alternatively activated M2. However, a more precise definition places them along a spectrum of activation where they may exhibit a number of pro- or anti-inflammatory roles. To gain a greater understanding of the mechanisms of the immune response from macrophages and the balance between M1 and M2 activation, we utilized two different modeling techniques, ordinary differential equation (ODE) modeling and agent-based modeling (ABM), to simulate the spectrum of macrophage activation to general pro- and anti-inflammatory stimuli on an individual and multi-cell level. The ODE model includes two hallmark pro- and anti-inflammatory signaling pathways and the ABM incorporates similar M1-M2 dynamics but in a spatio-temporal platform. Both models link molecular signaling with cellular-level dynamics. We then performed simulations with various initial conditions to replicate different experimental setups. Similar results were observed in both models after tuning to a common calibrating experiment. Comparing the two models’ results sheds light on the important features of each modeling approach. When more data is available these features can be considered when choosing techniques to best fit the needs of the modeler and application.

## Introduction

Macrophage polarization refers to the approximate state of activation of a macrophage responding to its environment. Macrophages show high plasticity and result in heterogenic subpopulations or polarized states identified by specific cellular markers [1]. Macrophage phenotypes may be largely classified as either pro-inflammatory or pro-injurious, also called classical macrophage polarization, or they can reflect an alternative activation profile, which has been considered as anti-inflammatory or pro-repair [2–4]. Classically activated (M1) macrophages promote the development of acute injury, whereas alternatively-activated macrophages (M2) may be involved in limiting or resolving inflammation [1]. Macrophage polarization is highly involved in physiological transitions from inflammation to tissue regeneration. A major field of macrophage biology seeks to understand the mechanisms and pathways leading to macrophage polarization. Macrophage polarization depends heavily on the tissue microenvironment and disease or injury state in which the cells are responding. Computational models provide an avenue to examine the many variables leading to macrophage polarization states.

The plasticity of macrophages has a significant impact on the overall ability of the immune system to resolve the insult [5]. Several mathematical models have been published that include macrophage polarization, including ODE models of subcellular signaling and simplified M1/M2 activation. Maiti et al. [6] and Moya et al. [7] focused on the subcellular signaling pathways of NF-*κ*B/TNF*α* and STAT3/IL-10, respectively. Frank et al. [8] and Zhao et al. [9] developed two-dimensional ODE models with M1 and M2 activation as the state variables. Rex et al. [10] used a Boolean model to select genes related to M1/M2 dynamics and developed an ODE modeling the dynamics of those genes. Additionally, some modeling efforts of macrophage plasticity incorporate spatial dynamics. Agent-based models that include M1/M2 phenotypes have been developed in the context of tuberculosis Kirschner et al. [11] and Nickaeen et al. [12] developed a PDE model of M1/M2 macrophages in response to high levels of IL4 or LPS/IFN*γ*.

In this work, we propose two models of the immune response to general inflammation that build upon previous models to examine the spectrum of macrophage activation in greater detail [3,13,14]. This model is an extension of work by Maiti et al. [6]; we added details of the IL-10 pathway not yet included in Maiti et al. by adapting and extending equations from Moya et al. [7], including both pro- and anti-inflammatory feedback loops and their interactions. The model consists of ten macrophages, each of which has a set of equations modeling its subcellular pathways. These ten macrophages are linked by external TNF*α* and IL-10, which can be both introduced into the system at various times and produced by the macrophages themselves. In our ABM, we incorporated pro- and anti-inflammatory mediators, allowed for M1/M2 activation to occur on a spectrum, and accounted for spatial dynamics. In this model, macrophages can become more activated towards an M1 or M2 phenotype based on their local patch environment, and perform a variety of roles depending on their activation levels. Both models account for macrophage cell cycle using randomly generated lifespans for each macrophage.

Based on data from Maiti et al. [6], we calibrated the models to each other by simulating a single macrophage with both pro- and anti-inflammatory stimuli. Through this initial scenario, we found that modeling the SOCS regulatory feedback loop is important in the definitive resolution of inflammation. We then simulated additional scenarios highlighting the effects of incorporating cell lifespan, recruitment, and various types of external stimuli and initial conditions. Comparison of these scenarios between the ODE model and ABM revealed overall similar behavior of M1 and M2 activation across two very different modeling approaches, suggesting that detailed subcellular pathway modeling is not necessary to achieve complex interplay between M1 and M2 polarization.

In the following sections, we describe the models in detail, the calibrating experiment, and the comparison of various simulated scenarios.

## Methods

### ODE subcellular macrophage model

#### Biological summary

There are several main interactions involved in cell signaling pathways that we include in our model. First, extracellular signals such as TNF*α* and IL-10 bind to and unbind from their receptors on the cell surface. Receptors transmit signals to other proteins within the cell, which may become activated or phosphorylated [15]. These complexes induce activation of transcription factors, proteins that are responsible for translocating to the nucleus, where they control the transcription of specific genes in the DNA into mRNA. mRNA then undergoes translation in the cytosol, where the protein corresponding to the gene is assembled according to the mRNA sequence [16]. We also account for degradation of various components. We model this process using the law of mass action unless otherwise specified. Details for these interactions are given in the following sections.

TNF*α* triggers a signaling pathway that leads to activation of the transcription factor NF*α*B and the subsequent shift to an M1 phenotype [17]. This results in the production of additional TNF*α* and IL-10 as well as other proteins. Alternatively, IL-10 activates the transcription factor STAT3 through the Jak-STAT pathway, giving rise to M2-type activation [18]. To capture the interactions between these pathways, we developed an ODE model, adapted from Maiti et al. [6] that includes these hallmark signaling pathways. This involves subcellular interactions between receptors and proteins in the cytosol and nucleus of the macrophage.

The model by Maiti et al. [6] initiates their signaling cascade with LPS, a molecule found in Gram-negative bacteria used to experimentally induce an immune response. IKK, a protein whose role is to regulate phosphorylation of I*κ*B*α*, is activated by both LPS and TNF*α*. Since we model general lung inflammation rather than a bacterial infection, we do not rely on activation of the M1 pathway by LPS; rather, we focus on activation via TNF*α*. Maiti et al. [6] include production of IL-10 and STAT3; our model extends this by including additional components of the Jak-STAT pathway and the negative feedback loops required to resolve the immune response. In the following sections, we note specifically which equations and terms are novel to our model.

LPS binds to its receptor, TLR4, which activates neutral IKK. IKK then phosphorylates I*κ*B*α* in the I*κ*B*α*-NF*κ*B complex, freeing NF*κ*B to translocate to the nucleus. In the absence of a stimulus, I*κ*B*α* sequesters NF*κ*B to prevent it from causing the production of unnecessary proteins. Transcription factor NF*κ*B initiates transcription of TNF*α*, IL-10, A20, and I*κ*B*α* mRNA, resulting in their translation and protein production. As part of a negative feedback loop that prevents excessive production of these proteins, A20 inactivates active IKK and I*κ*B*α* sequesters unbound NF*κ*B. TNF*α* and IL-10 are secreted from the cell.

Extracellular TNF*α* binds to its receptor, activating neutral IKK. IL-10 also binds to its receptor, and JAK and Tyk tyrosine kinases, whose main function is to activate STAT3, bind to this complex as well. Without all of these components, STAT3 cannot be phosphorylated and control transcription of key genes in the nucleus. The IL-10-Jak-Tyk complex activates STAT3, which translocates to the nucleus and initiates the production of IL-10, SOCS1, and SOCS3. Both SOCS1 and SOCS3 are part of negative feedback loops that bring about resolution of both the M1 and M2 pathways. SOCS3 inhibits transcription of TNF*α* mRNA and both SOCS1 and SOCS3 inhibit activation of STAT3. IL-10 also inhibits activation of IKK.

Eqs (2) through (27) are from Maiti et al. [6] unless otherwise noted, and the model variables we added are shown in Eqs (28) through (38). Fig 1 summarizes these interactions, described in more detail in the equations. This schematic describes interactions between receptors, transcription factors, and other proteins within the cell in response to detection of extracellular signals on the cell surface. Table 1 lists the parameters used in the model and their descriptions. Code for these equations can be found in the supplement.

**Fig 1.**
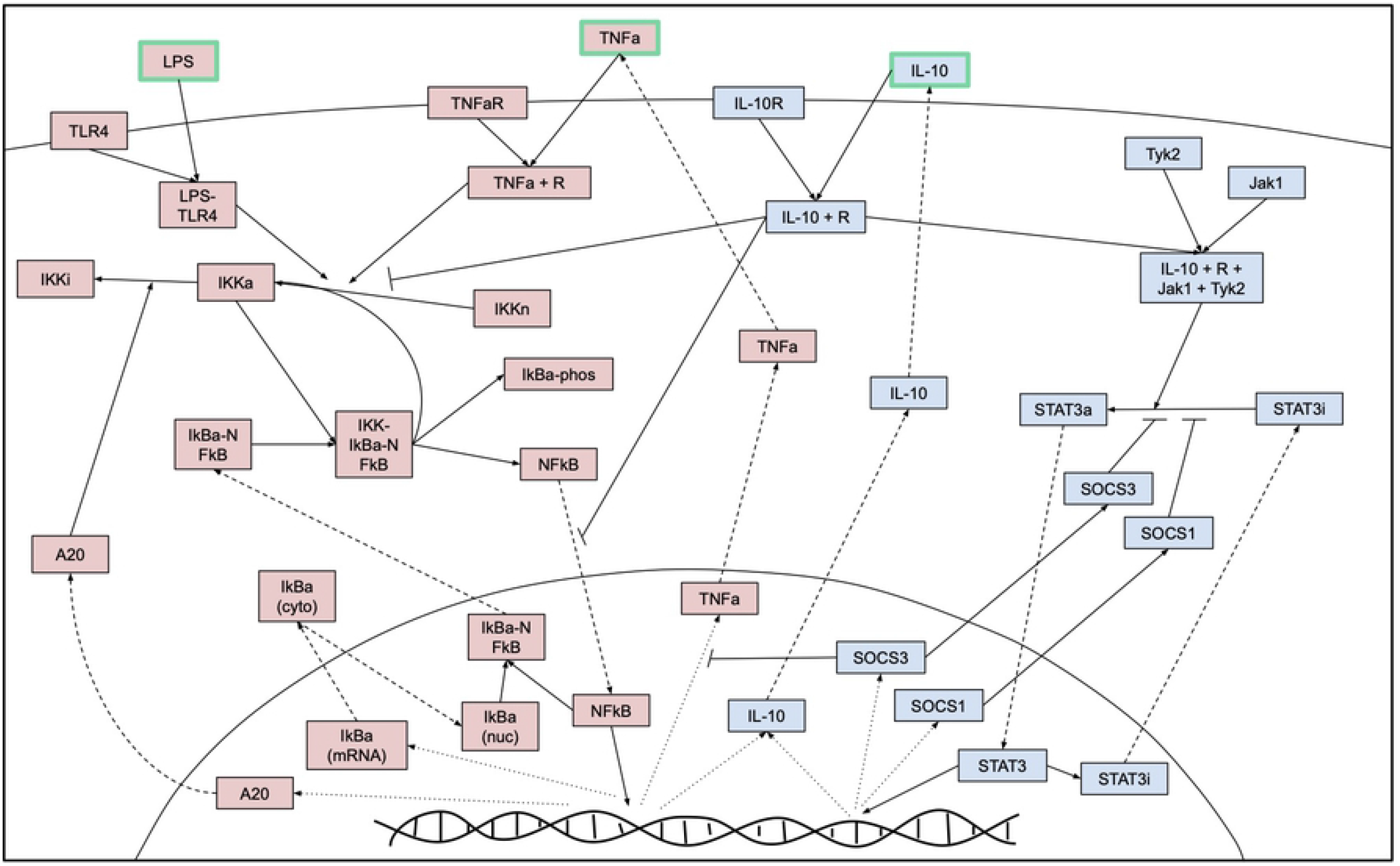
Model schematic. Schematic of interactions within a macrophage and with external stimuli LPS, TNF*α*, and IL-10. Dashed lines represent interactions that involve movement between the cytosol and nucleus, dotted lines represent transcription processes in the nucelus, and solid lines represent all other interactions. Red boxes represent components that are primarily associated with the pro-inflammatory/M1 pathway and blue boxes with the anti-inflammatory/M2 pathway.

**Table 1.**
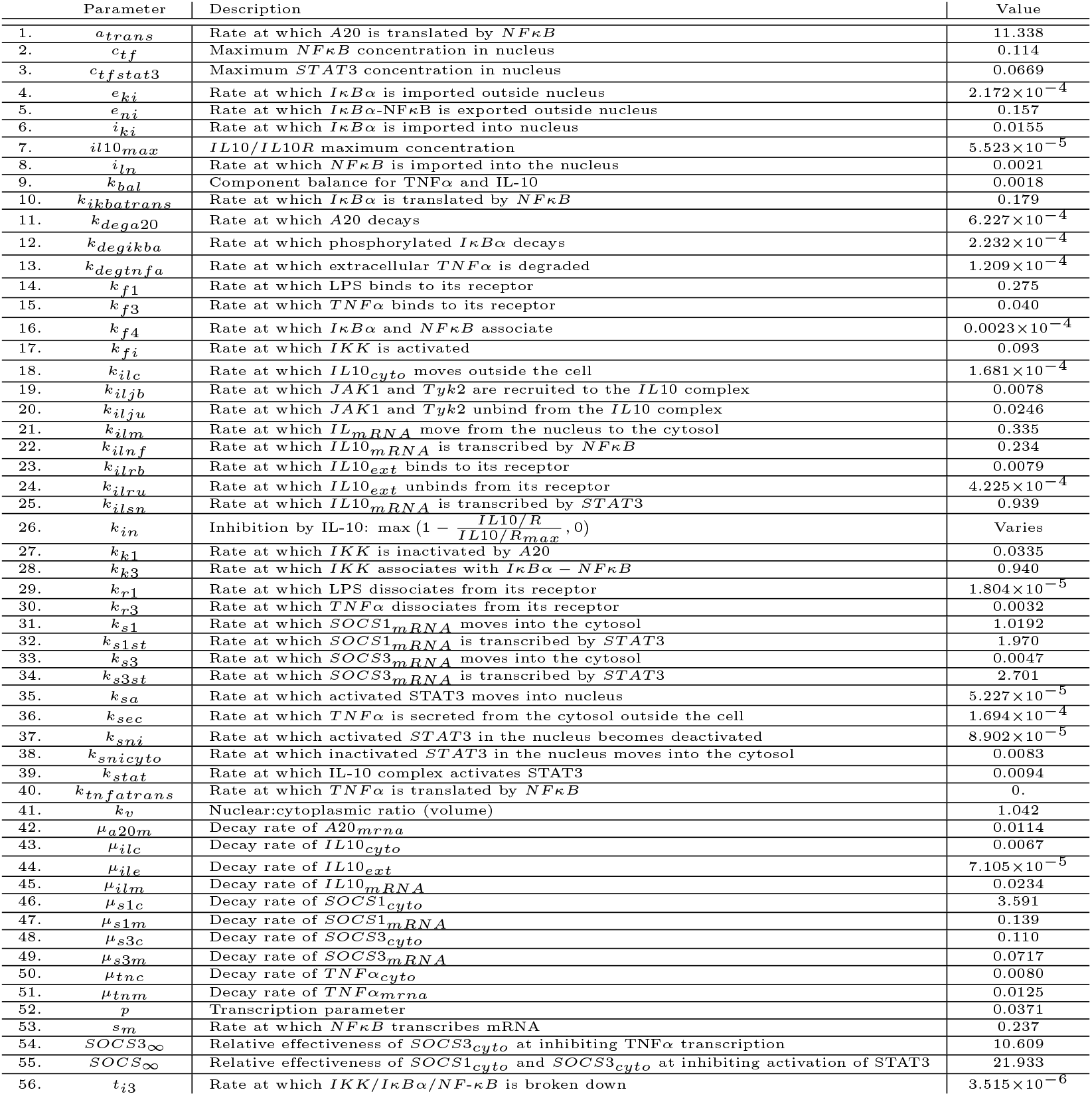
List of parameter estimates from preliminary fit for the subcellular pathways model.

#### LPS

Maiti et al. began the model through initiation by LPS, a major component of bacteria identified by the macrophage. LPS is represented as a constant input into the system, shown in Eq (1). When LPS is detected by TLR4, its receptor, they form a complex denoted *LPS/TLR4*, shown in Eqs (2) and (3). Components connected by a forward slash, such as *LPS/TLR4*, represent a complex; otherwise, variables side by side are multiplied together. We will use this convention in the equations described throughout this section.

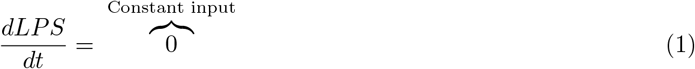

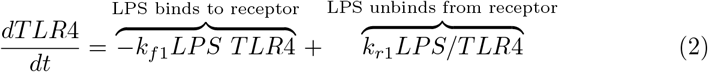

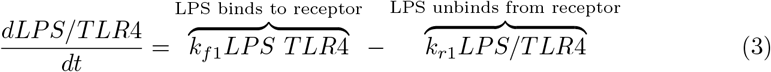

#### I*κ*B*α* kinase

I*κ*B*α* kinase (IKK) is represented in three distinct states: neutral, active, and inactive, shown in Eqs (4), (5), and (6), respectively. The binding of LPS and TNF*α* to their respective receptors triggers the activation of neutral IKK, represented by the first term in Eqs (4) and (5). As part of a negative feedback loop for the pro-inflammatory response, IL-10 inhibits neutral IKK from activating. Maiti et al. describes this inhibition in the first term of Eqs (5) and (5) through the parameter *k_in_*, where

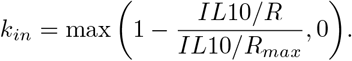

Active IKK phosphorylates the IKK-I*κ*B*α*-NF*κ*B complex (second term in Eq (5)). Phosphorylation causes the complex to break down, releasing a neutral form of IKK, shown in the second term of Eq (4). Finally as part a negative feedback loop to prevent an overactive pro-inflammatory response, the protein A20 inactivates active IKK, the last term of Eq (5) and Eq (6).

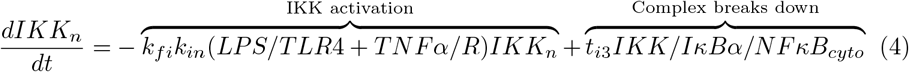

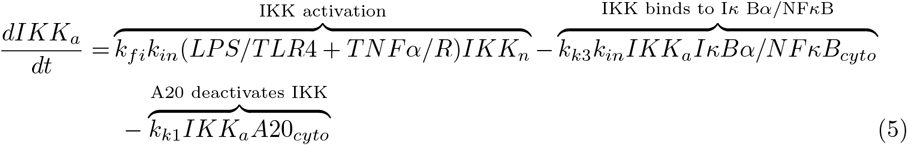

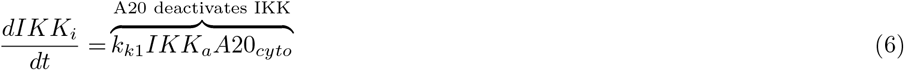

#### I*κ*B*α*

In a resting state, I*κ*B*α* sequesters free NF*κ*B by associating into a complex, shown in the first term of Eq (7). This process also occurs in the nucleus, from which the complex can move to the cytosol (second term of Eq (7)). Activated IKK phosphorylates the complex, represented by the third term in Eq (7). The binding of active IKK to I*κ*B*α*-NF*κ*B (first term of Eq (8)) causes all three components to separate, modeled by the second term of Eq (8): NF*κ*B is released, I*κ*B*α* is degraded, and IKK returns to a neutral state.

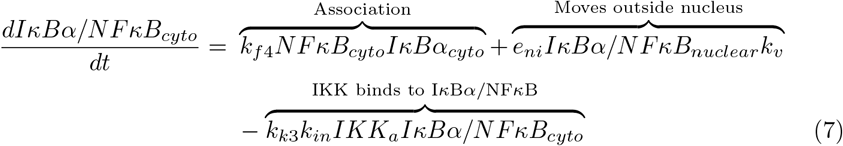

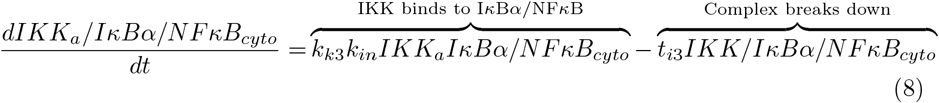

Eqs (9) through (12) show the various states of the inhibitory protein I*κ*B*α*. NF*κ*B promotes the transcription of I*κ*B*α* mRNA, shown in the first term of Eq (9). Subsequent translation of the protein and decay of the mRNA are described in the first term of Eq (10) and the second term of Eq (9), respectively. As previously described, the second term of Eq (10) represents I*κ*B*α* sequestering free NF*κ*B in the cytosol. In a resting cell, excess I*κ*B*α* is distributed evenly between the cytosol and nucleus; thus, the last two terms of Eq (10) show import and export of I*κ*B*α* between the two compartments [19]. The parameter *k_v_* accounts for the nuclear-cytoplasmic ratio to account for the size of the cell’s cytoplasm in relation to its nucleus. The release of NF-κB from the I*κ*B*α*-NF-*κ*B complex by active IKK results in the phosphorylation of I*κ*B*α* and its subsequent degradation, shown in the two terms of Eq (12).

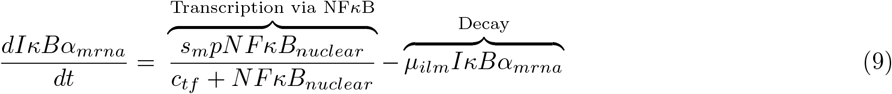

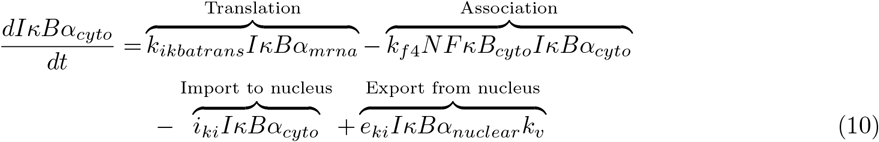

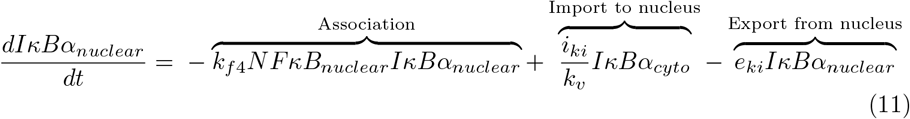

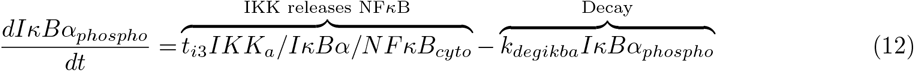

#### NF*κ*B

The protein NF*κ*B is released from the complex (first term of Eq (13)) and translocates to the nucleus, represented by the second term of Eq (13) [19]. NF*κ*B activates the transcription of several genes, including TNF*α* and IL-10, A20, and I*κ*B*α*. I*κ*B*α* sequesters nuclear NF*κ*B (last term in Eq (14) and first term in Eq (15)) before the complex moves back into the cytosol, shown in the last term of Eq (15).

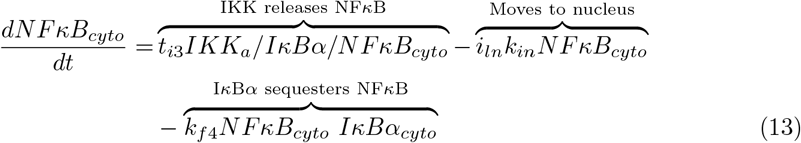

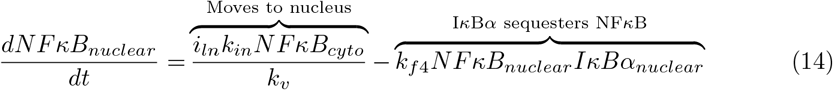

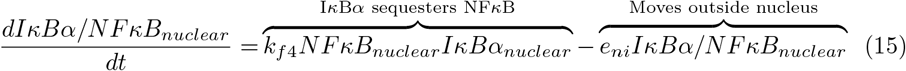

#### TNF*α*

One of the main targets of gene expression of NF*κ*B is the pro-inflammatory cytokine TNF*α*. The first term of Eq (16) represents transcription of mRNA. There is evidence that Suppressor of Cytokine Signaling 3 (SOCS3), discussed in further detail below, plays a role in regulating the pro-inflammatory response by inhibiting TNF*α* mRNA and protein production, although the exact mechanisms by which this occurs is still unclear [20,21]. We included a multiplier, not in the original equation by Maiti et al., in this first term to represent inhibition of mRNA production by SOCS3. After transcription and translation, TNF*α* is secreted from the cell (first two terms of Eq (17)). The parameter *k_bal_* represents a component balance for TNF*α* as it moves from the cytosol to the supernatant.

Extracellular TNF*α* binds to its receptor on the cell surface, represented by the second term in Eq (18). In some cases the cytokine unbinds from its receptor, accounted for by the second term in Eq (18). Once inside the cell, either after binding to its receptor or being translocated from the nucleus, TNF*α* performs several important roles. Shown in the first term of Eq (4), TNF*α* bound to its receptor upregulates activation of IKK, which then precipitates further NF*κ*B transcription.

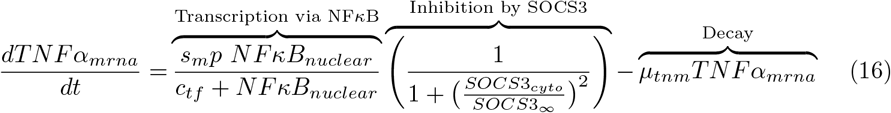

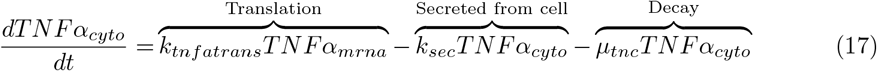

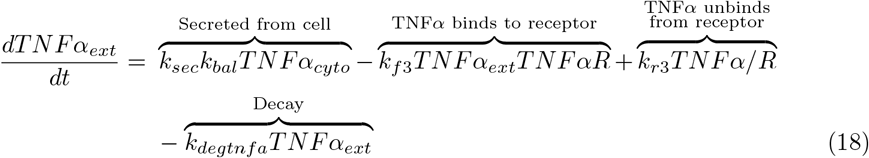

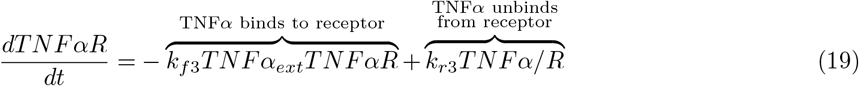

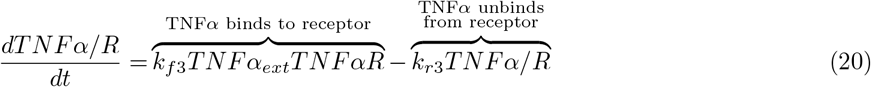

#### A20

As mentioned previously, A20 is another NFκB-responsive gene responsible for deactivating IKK, which blocks NF*κ*B translocation to the nucleus. Eq (21) shows transcription and subsequent degradation of A20 mRNA. Eq (22) shows translation of the protein in the cytosol, and A20 decays at rate *k*_*dega*20_, second term in Eq (22).

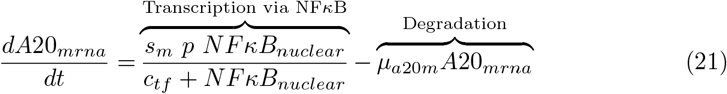

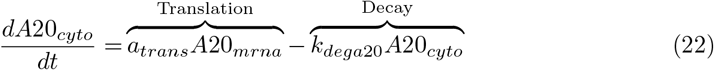

#### IL-10

A hallmark of the anti-inflammatory response is the cytokine IL-10. Its gene is a target of NF*κ*B transcription and is involved in the regulation of the pro-inflammatory response. Some events related to IL-10 production and function are included in the model by Maiti et al. [6], but we expand the model to include a fuller view of the role of IL-10 and an important pathway it activates.

Extracellular IL-10 can bind to and unbind from its receptor IL-10R, as modeled by the first two terms in Eq (23) [7]. For simplicity, we assume the total number of receptors is conserved. The first term in Eq (25) describes upregulation of the IL-10 gene by transcription factors NF*κ*B and STAT3. Maiti et al. include the constants 0.4 and 0.6 such that NF*κ*B is responsible for 40% of the transcription rate and STAT3 is responsible for the other 60%. The nonlinear terms represent maximum possible rates of IL-10 transcription, since space in the nucleus is limited. IL-10 is translated from its mRNA and secreted from the cell (first two terms of Eq (26)). The third term in Eq (23) includes a component balance *k_bal_* between the cytosol and supernatant. Baseline degradation rates for extra- and intracellular IL-10 and IL-10 mRNA is included in Eqs (23), (26), and (25), respectively.

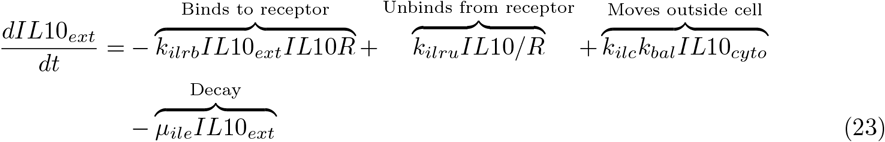

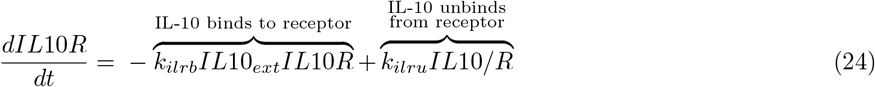

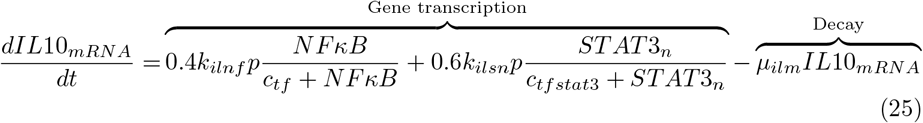

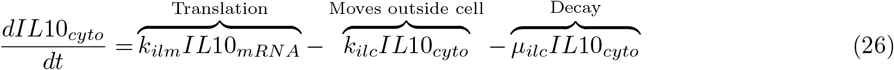

#### JAK-STAT signaling

Aside from inhibitory functions, IL-10 signaling initiates the JAK-STAT signaling pathway, a primary mechanism through which the immune response mediates inflammation [22]. The protein tyrosine kinases JAK1 and Tyk2 are recruited to the IL-10/IL-10 receptor complex, shown in the third term of Eq (27). This creates a new complex, *IL*10/R/*JAK*1/*Tyk*2, Eq (30) [23]. The second term accounts for the possibility that the complex may break apart. JAK1 (Eq (28)) and Tyk2 (Eq (29)) concentrations are conserved, assuming enzyme-type dynamics. In light of the many components involved in creating this complex, we explored incorporating the various combinations of the binding steps, such as the individual receptor components, each of which bind to a specific tyrosine kinase. In the end, we decided to model the recruitment of JAK1 and Tyk2 to the IL-10/IL-10 receptor complex as one step; this still captures the appropriate dynamics without adding more parameters and equations. The last two terms of Eq (27) and all of Eqs (28) through (30) are our additions to the original model by Maiti et al., with terms representing activation of STAT3 through the Jak-STAT pathway adapted from Moya et al. [7].

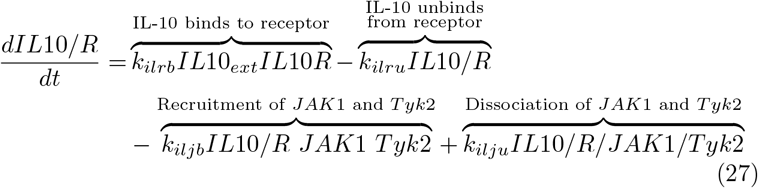

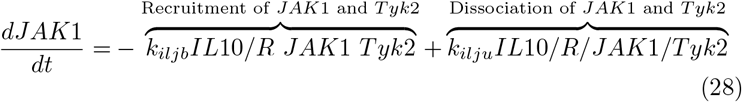

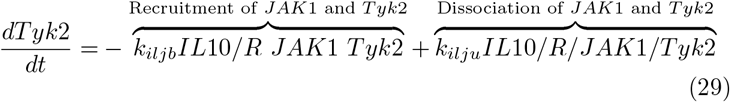

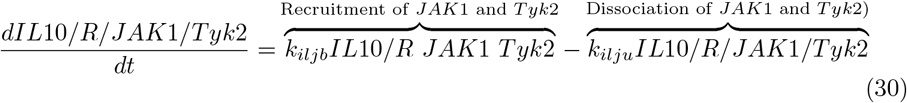

The IL-10/IL-10 receptor/JAK1/Tyk2 complex serves as a temporary docking station for inactive Signal Transducer and Activator of Transcription 3 (STAT3) [24]. Upon recruitment to the complex, STAT3 is activated and undergoes homodimerization, shown in the first term of Eq (31). Maiti et al. modeled the recruitment and activation of STAT3 through binding of STAT3 to the IL-10/IL-10R complex without Jak1 and Tyk2. We also included a multiplier representing inhibition by Suppressors of Cytokine Signaling 1 and 3 (SOCS1 and SOCS3), two IL-10 responsive genes as well as the second term of Eq (33) and Eq (34) which allow for the conservation of STAT3 in the model. SOCS1 inhibits JAK1 function by binding its SH2 domain to JAK1, preventing STAT3 from docking to the IL-10 complex. SOCS3 performs a similar role but docks to the receptor; since we do not model at the level of detail of specific binding locations, we model this inhibition as having the same result, which is preventing STAT3 from activating [25–27].

STAT3 translocates to the nucleus (second term of Eq (32)) and controls transcription of several IL-10 responsive genes. The main inhibitor of STAT3 function is PIAS3. The protein binds to activated STAT3, preventing further transcription [28]. We model this by including a deactivation term with rate *k_sni_*, shown in the second term of Eq (33). Assuming enyzme-type dynamics for all states of STAT3, the transcription factor is conserved, and deactivated nuclear STAT3 returns to the cytosol in the last term of Eq (34).

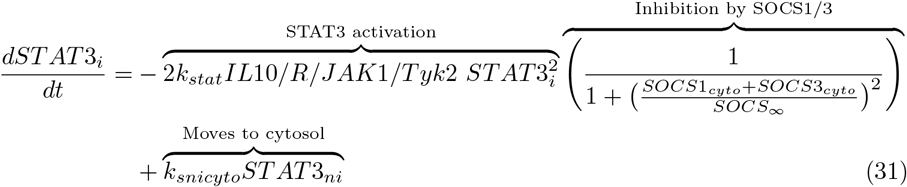

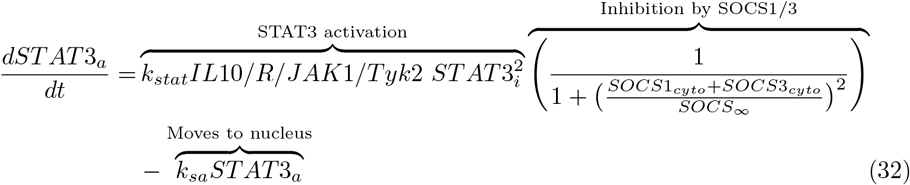

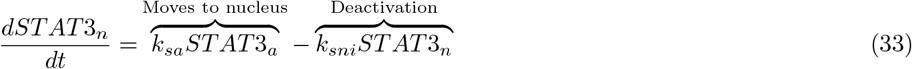

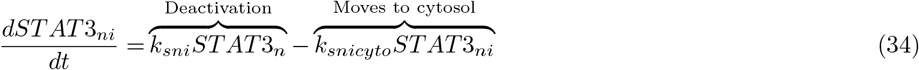

#### SOCS

The inclusion of SOCS, represented in Eqs (35) through (38), is also novel to our model as compared to that by Maiti et al. Suppressors of Cytokine Signaling 1 and 3 (SOCS1, SOCS3) are upregulated via STAT3 transcription and translation, first two terms of Eqs (35) and (36), respectively [18, 29]. The last terms of these two equations represent natural degradation of the mRNA.

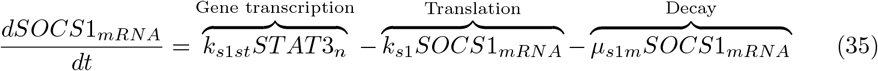

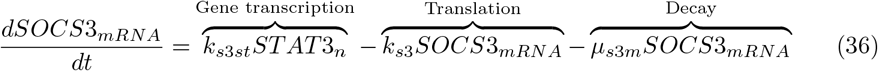

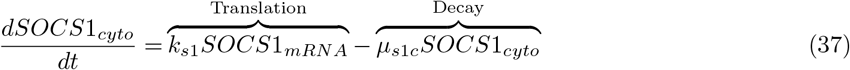

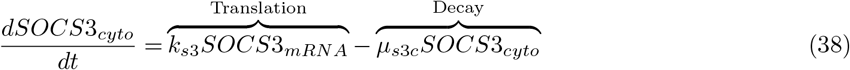

We used this model that includes both pro- and anti-inflammatory signaling pathways to provide a fuller picture of the spectrum of activation that can occur within a macrophage. In the following pages, we discuss how this model was implemented and compared to the ABM.

### Parameters & initial conditions for ODE model

We used data from Maiti et al. [6] to obtain initial parameter values as a starting point. This was not a complete parameter estimation (calculating sensitivities, etc.), but rather a first step in obtaining parameter values and initial conditions that produce dynamics that are roughly expected and have the correct scales. The data provided by the authors was processed data and did not align exactly with our model variables, so we used their parameters as a starting point and adjusted to get similar fits to their dynamics.

Since the model simulations by Maiti et al. [6] were initialized with LPS, once the final parameter set was obtained the model was run for 1,000 hours with no LPS using the code provided in the supplement. The ending values of these simulations for each variable were determined to be the baseline initial conditions, representing a state of no macrophage activation.

### Modeling multiple macrophages

The equations described in the section above represent the pathways in a single macrophage. To model recruitment and cell lifespan, we extended the model to represent ten identical macrophages. These macrophages share the same extracellular components: LPS, IL-10, and TNF*α*. Figure 2 shows a visualization of this compartmental model. Furthermore, each macrophage is randomly assigned a lifespan, 12 ± 3 hours. At the end of each cell’s lifespan, the variables in the signaling pathway are returned to a naive state to represent the recruitment of a naive cell.

**Fig 2.**
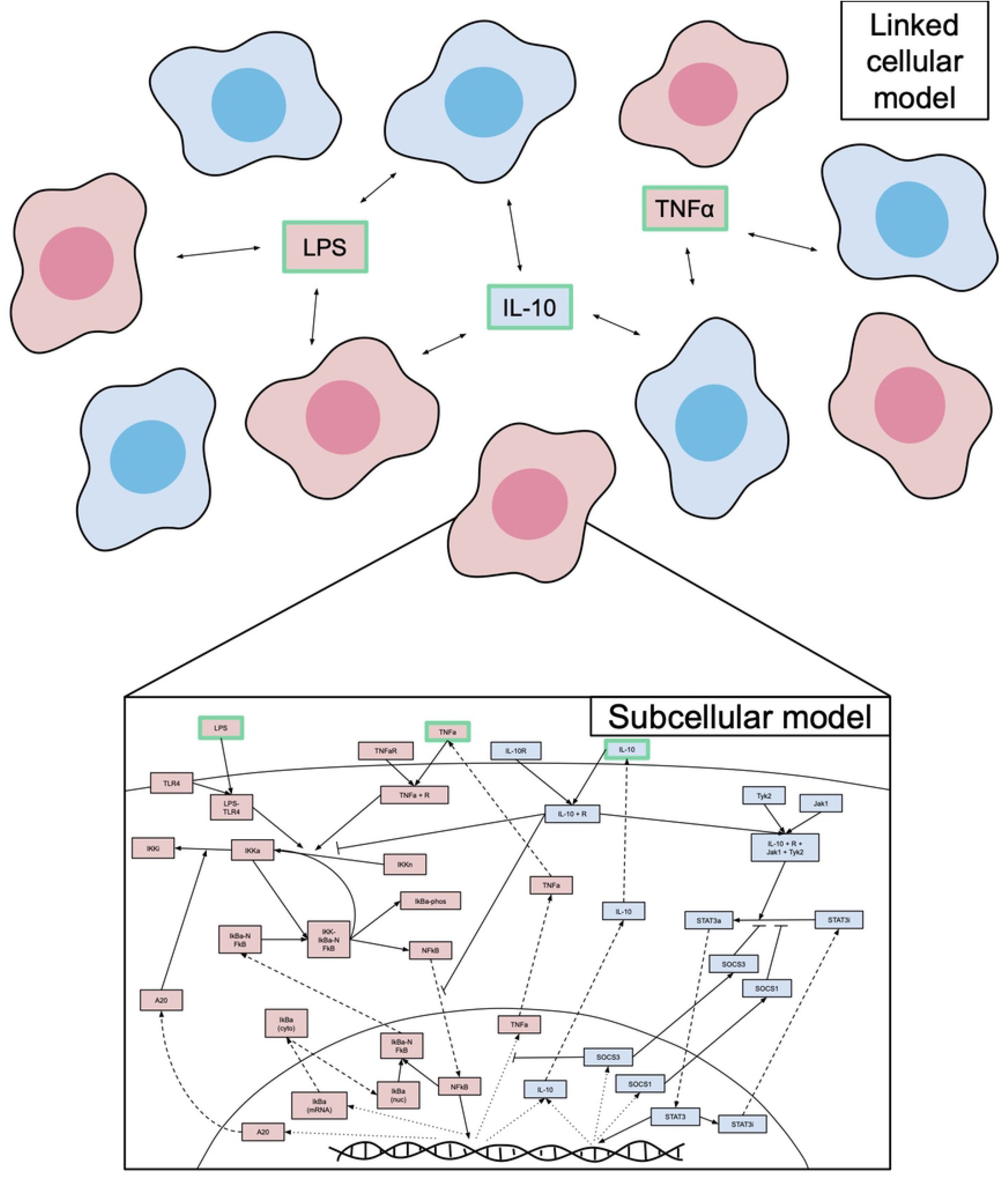
Cellular and subcellular scales in the model are linked by extracellular signals. A representation of the multiple macrophages ODE model, in which each macrophage is a compartment with its own set of subcellular signaling pathways, and all ten macrophages share external stimuli LPS, TNF*α*, and IL-10.

Our aim in constructing a model of multiple macrophages was to examine how macrophages in close proximity behave in response to extracellular stimuli while still utilizing the ODE structure. Resulting dynamics of variables that exist in each model can be viewed separately or averaged together to obtain the average behavior across all macrophages.

### Agent-based M1/M2 model

Our ABM tracks pro- and anti-inflammatory mediators (PIM and AIM, respectively), M0, M1, and M2 macrophages, and SOCS on a 40-by-40 grid, implemented using object-oriented programming in MATLAB (code provided in the supplement). Macrophages are mobile agents with M1/M2 activation and SOCS levels as associated attributes. Each macrophage may take up one patch, and pro- and anti-inflammatory mediators are measured by amount on each patch, diffusing across the grid over time. We do not specifically model particular cytokines, but rather the general levels of pro- and anti-inflammatory mediators. The model can be initialized with varying levels of any of these components and simulated to obtain the resulting dynamics. The model performs a series of steps to recruit macrophages, determine M1/M2 activation, and produce and inhibit pro- and anti-inflammatory mediators and SOCS. Each macrophage has levels of M1 and M2 activation, where 0 ≤ M1 + M2 ≤ 1, and these activation levels are updated based on the surrounding levels of pro- and anti-inflammatory mediators. Figure 3 summarizes the steps taken during every iteration of the simulation, where each iteration represents 20 minutes. These steps are based on the same interactions described in the ODE model.

**Fig 3.**
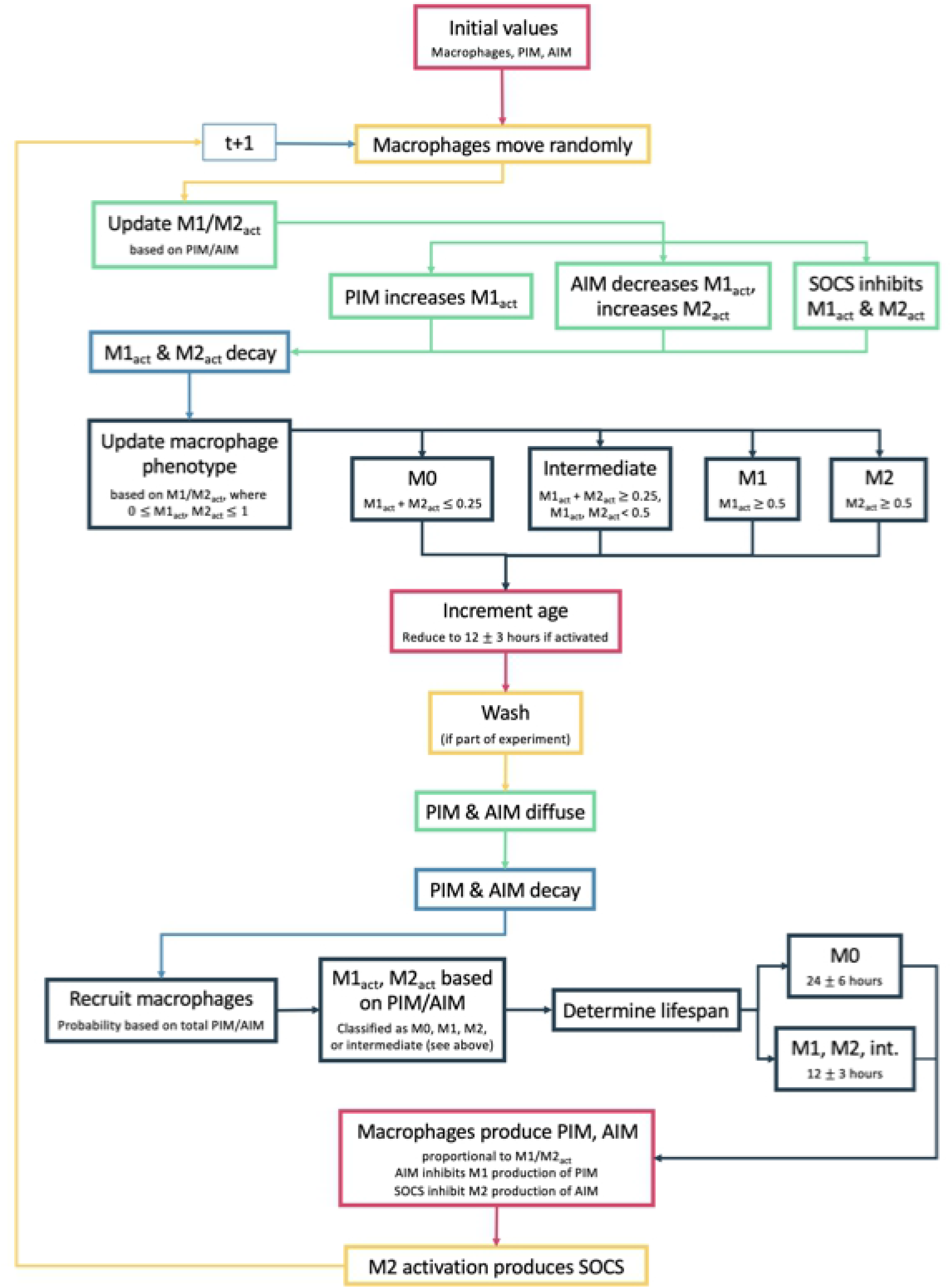
Flow chart for ABM rules. Description of steps in ABM for each iteration of the simulation.

### Calibrating experiment and scenarios

To compare the models to each other, we implemented the same scenario, which we called a “calibrating experiment,” in each model and tuned the ABM results so that PIM & AIM and M1 & M2 activation results were similar to their corresponding components in the ODE model, since the ODE model parameters were already set (see section describing ODE model parameters for process). These ODE model components were extracellular TNF*α* & IL-10 and TNF*α* mRNA & IL-10 mRNA, respectively. We chose M1 and M2 activation to be represented by TNF*α* and IL-10 mRNA, respectively, since mRNA is produced via downstream signaling initiated by the surrounding environment and also results in specific proteins that are secreted from the cell. Thus, mRNA associated with the cell’s phenotype both reflects and drives macrophage polarization.

Tuning parameters so that the ABM and ODE model returned similar dynamics in the calibrating experiment allowed us to obtain similar behavior at baseline and compare the results of more complicated experiments. We chose this scenario to be a single macrophage with a high pro-inflammatory stimulus and without cell death. In the ODEs, initial conditions were established such that all variables are at baseline levels, to represent an M0 macrophage. TNF*α*, the variable representing a pro-inflammatory stimulus in the ODE model, was set to 10 pg/mL, consistent with experimental methods [30].

For a naive macrophage in the ABM, we used a 3-by-3 grid so that the cell could move but interact only with the mediators in its immediate proximity. A naive macrophage in this model is defined as having activation M1 + M2 < 0.25; M1 and M2 activation were randomly chosen with bounds that satisfy this condition. Pro-inflammatory mediators do not have specific units but after exploratory simulations, we considered a concentration of 30 in the center space of the grid to be sufficient to mount an inflammatory response. ABM parameters were tuned manually to match the dynamics observed in the ODE.

Through simulating the calibrating experiment and scenarios, described below, we found that receptor-bound TNF*α* and IL-10 in the ODE model played an important role in the resulting dynamics. Many modelers do not model changes in cytokine levels due to binding to receptors, assuming this amount is negligible. However, we found that this is not the case in our ODE model, and explicitly modeling receptors makes a difference in dynamics. Receptors were not explicitly modeled in the ABM; macrophage activation is based solely on the surrounding PIM and AIM. This can create a disparity in the amount of PIM and AIM that are compared between the two models. In Figures 4 and 5 and in our results, we showed two cases of the ODE model: when only extracellular TNF*α* and IL-10 are considered, and when both extracellular and receptor-bound TNF*α* and IL-10 are considered. We also discussed differences between these two cases.

**Fig 4.**
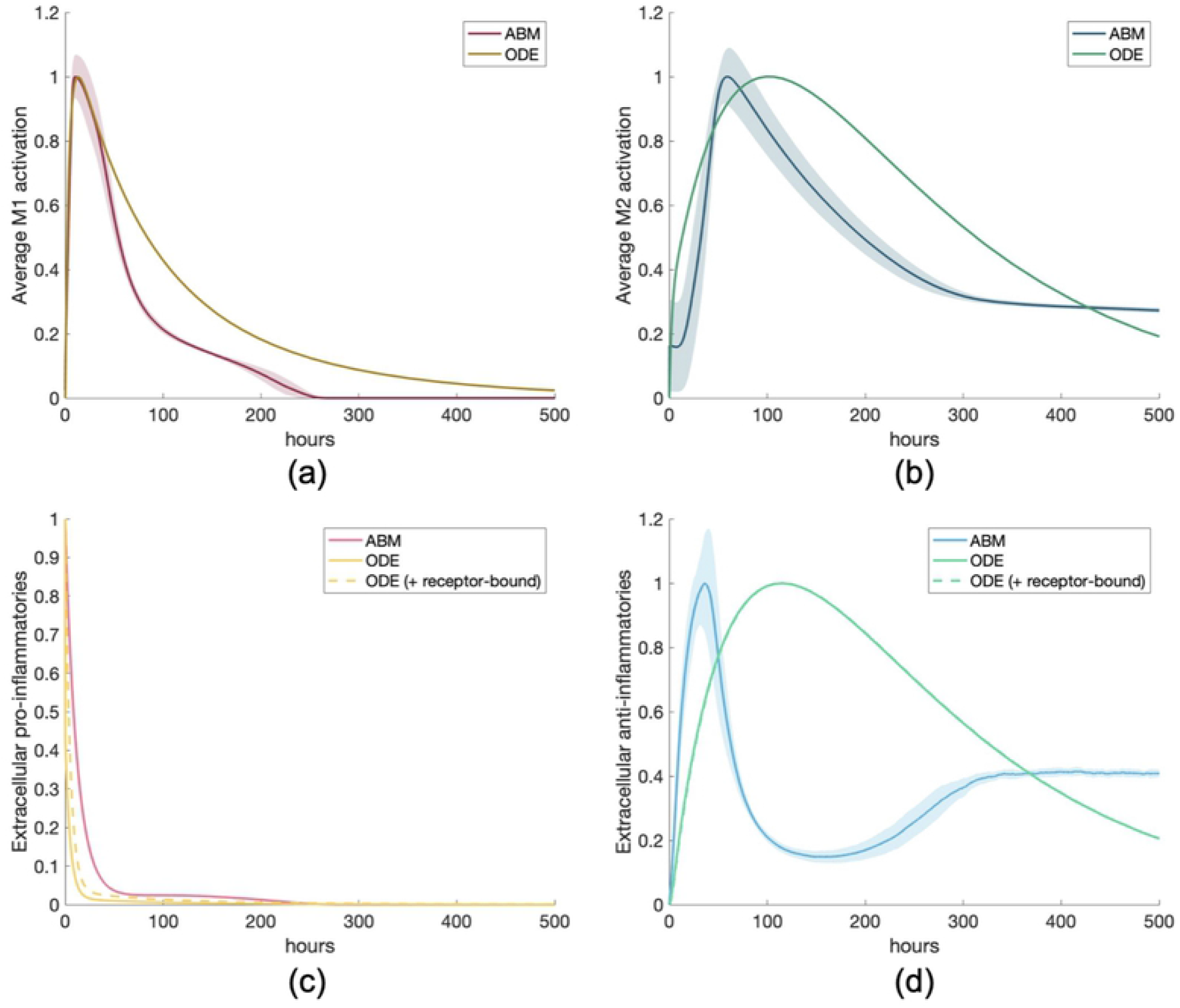
Results for calibrating experiment show good agreement between models. Calibrating experiment: single macrophage activated by a pro-inflammatory stimulus. ABM and ODE results are shown on the same plots for comparison. All transients are scaled by their maximums. Dotted lines represent extracellular TNF*α* or IL-10 with receptor-bound TNF*α* or IL-10, respectively. (a) M1 activation, (b) M2 activation, (c) pro-inflammatory mediators, (d) anti-inflammatory mediators.

**Fig 5.**
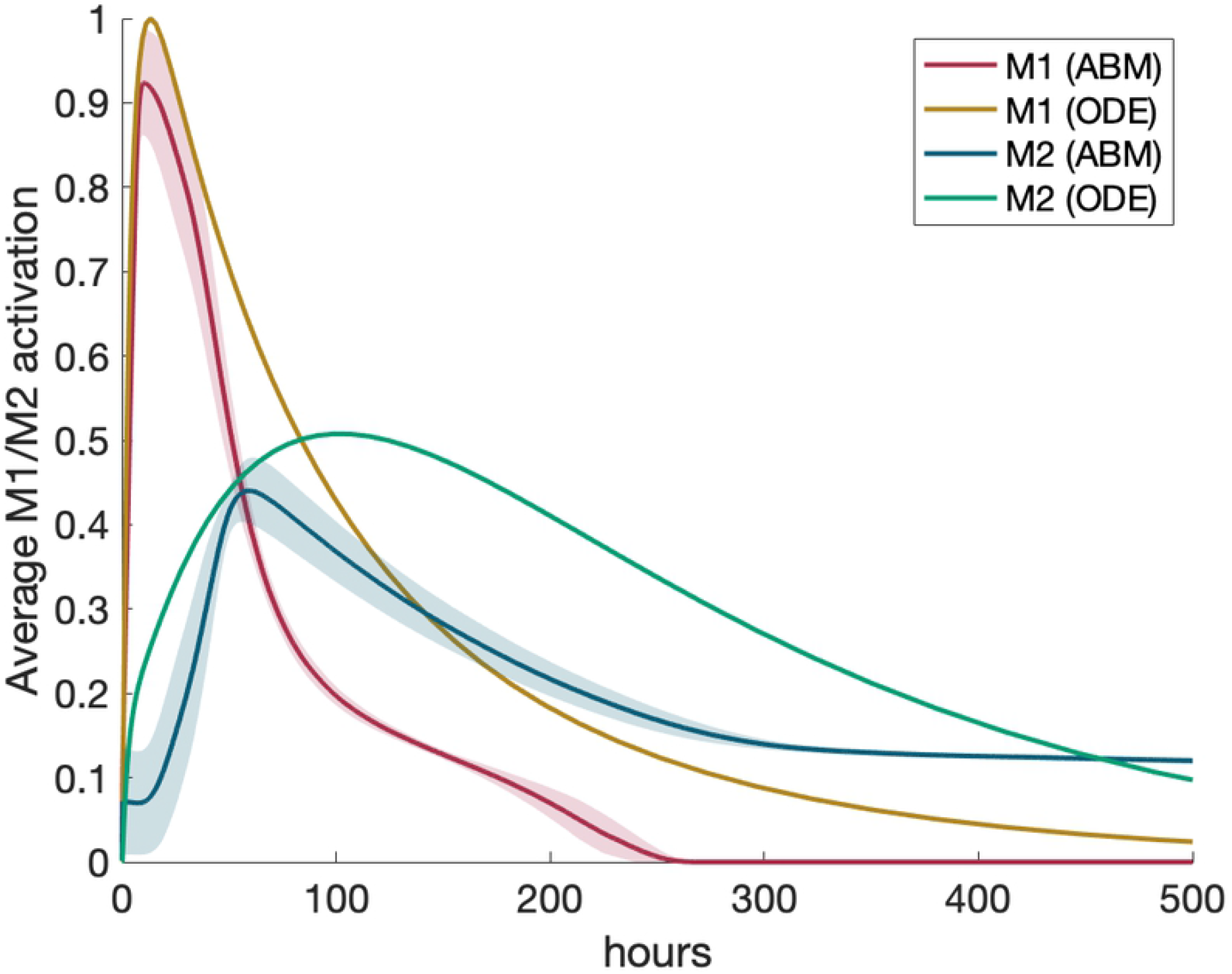
Results for M1 activation relative to M2 activation show good agreement between models. Calibrating experiment: M1 and M2 activation resulting from the calibrating experiment. ODE results are scaled by the maximum M1 activation to compare to activation in the ABM, which is bound by 0 and 1.

In the calibrating experiment, we set the ODE and ABM parameters and initial conditions such that a single macrophage would exhibit similar M1 and M2 behavior when initialized with PIM (process described in Methods section). Figure 4 shows the results of this simulation. All transients are normalized for comparison because the units in the models vary. To do this, we scaled each transient by its maximum. The results of the ABM in Figure 4 is the result of 50 simulations; on the other hand, the ODE model with a single macrophage is deterministic and thus only one simulation is necessary.

M2 activation occurs slightly earlier in the ABM than in the ODE, but we concluded that the results were similar enough to proceed with comparisons. Adding receptor-bound TNF*α* and IL-10 to their extracellular counterparts did not make a significant difference in the results. We also considered the magnitudes of M1 and M2 activation in relation to each other, shown in Figure 5. M1 and M2 activation in the ABM are, by definition, bound between 0 and 1. To compare with the ABM, we scaled TNF*α* and IL-10 mRNA in the ODE by the maximum of TNF*α*. Peak M2 activation in both the ABM and ODE are about half the peak M1 activation. This shows important dynamics observed in both models, which illustrates the strength of the calibration.

Once the parameters were set and the calibrating experiment was simulated, we changed the initial conditions to represent six additional scenarios, which will be described in greater detail below. First, we used the same single-macrophage model as described above but with an anti-inflammatory stimulus. Then, using the 40-by-40 grid for the ABM and ten-macrophage model for the ODE, we incorporated recruitment/turnover and cell lifespan. For these larger models, we simulated the following scenarios, the results of which will be discussed in the following section:

1. Naive macrophages with large pro-inflammatory stimulus
2. Naive macrophages with large anti-inflammatory stimulus
3. M1 macrophages with anti-inflammatory stimulus
4. Half M1, half M2 macrophages
5. Pro-inflammatory stimulus, wash at hour 12, then anti-inflammatory stimulus

### Sensitivity analysis

We performed a one-at-a-time sensitivity analysis (SA) on both the ABM and ODE models to identify the key parameters that drive macrophage activation and expression of pro- and anti-inflammatory mediators. Using the single-macrophage calibrating experiment, we perturbed each parameter (except for those related to lifespan) for both models by 10% above or below the original value and calculated the percent change for key variables and multiple time points. Parameters and their original values are shown in Tables 1 and 2 for the ODE model and ABM, respectively.

**Table 2.** ABM parameters and descriptions.

| Parameter | Description |
| --- | --- |
| ImmuneProInflammatoryRate | Rate at which M1s produce PIM |
| ImmuneAntiInflammatoryRate | Rate at which M2s produce AIM |
| ImmuneM1AntiInflammatoryRate | Rate at which M1s produce AIM |
| ProInflammatoryDecayRate | Rate at which PIM decay |
| AntiInflammatoryDecayRate | Rate at which AIM decay |
| SOCSDecayRate | Rate at which SOCS decay |
| PIMNegativeFeedbackRate | Rate at which M1 activation decays |
| AIMNegativeFeedbackRate | Rate at which M2 activation decays |
| RecruitmentMMTerm | Regulates effectiveness of macrophage recruitment by PIM & AIM |
| AIMRecruitScale | Rate at which AIM recruit macrophages |
| PIMActivationScale | Regulates effectiveness of M1 activation of newly recruited cells by PIM |
| AIMActivationScale | Regulates effectiveness of M2 activation of newly recruited cells by AIM |
| AIMInfinity | Regulates effectiveness of AIM in inhibiting M1 activation of newly recruited cells by PIM |
| M1ActivationRate | Rate at which M1 activation is increased by PIM |
| M1ActHillParameter | Regulates effectiveness of increasing M1 activation via PIM |
| M2ActScalar | Rate at which M2 activation is increased by AIM |
| M2ActHillParameter | Regulates effectiveness of increasing M2 activation via AIM |
| M1AIMInfinity | Regulates effectiveness of AIM in inhibiting M1 activation of local cells by PIM |
| M1DecreaseViaAIM | Rate at which M1 activation is decreased by AIM |
| M1DecreaseViaAIMHill | Regulates effectiveness of decreasing M1 activation by AIM |
| SOCSProductionRate | Rate at which AIM produce SOCS |
| AIMSOCSHill | Regulates effectiveness of SOCS production by AIM |
| M1SOCSInfinity | Regulates effectiveness of inhibiting M1 activation by SOCS |
| M2SOCSInfinity | Regulates effectiveness of inhibiting M2 activation by SOCS |
| AIMSOCSInfinity | Regulates effectiveness of inhibiting M2 production of AIM production by SOCS |

In this work, we are most interested in M1/M2 activation and pro- and anti-inflammatory mediators; thus, we track results for the corresponding model variables. Since lifespan is not considered, TNF*α*-induced dynamics persist for the four variables for about 1000 hours (see Figure 4). Thus, we obtained SA results for the following time points: 50, 100, 250, 500, 750, and 1000 hours.

## Results

We simulated equivalent scenarios in an ODE model and an agent-based model of M1/M2 activation in response to general inflammatory stimuli. In this section we compare the results of the two models to shed light on the benefits of each model type and, in particular, examine whether the incorporation of a spatial component through an ABM or the incorporation of hallmark signaling pathways through an ODE improve the value of the models in understanding immune system dynamics. All results shown are the average of 50 simulations except the single-macrophage simulations, which is deterministic since age is not a factor. Code is provided in the supplement.

### Scenario 1: Macrophage with anti-inflammatory stimulus

For the first scenario, we used the same structure of a single macrophage as in the calibrating experiment. Instead of a pro-inflammatory stimulus, we used an anti-inflammatory stimulus. Figure 6 shows the results of this simulation. AIM and M2 activation behave roughly the same; in the ABM, M2 activation decreases slightly slower than in the ODE. For the ABM, in both the calibrating experiment and this scenario, there is a slight increase in AIM later in time. This may be due to the small amount of SOCS left at this time, allowing AIM to increase slightly before decaying completely due to decreasing M2 activation. Including receptor-bound mediators in the ODE reveals a slower decrease in AIM over time but overall similar behavior to the ABM. A small increase in M1 activation and PIM also occurs later in time in the ODE model; this is due to trace amounts of NF*κ*B in the baseline levels of the cell that result in a small amount of TNF*α* production downstream. Additionally, we noted that simulating a single macrophage in the ABM shows consistent results for each of the 50 simulations, since the shaded regions around the curves, representing standard deviation, are very small or nearly zero.

**Fig 6.**
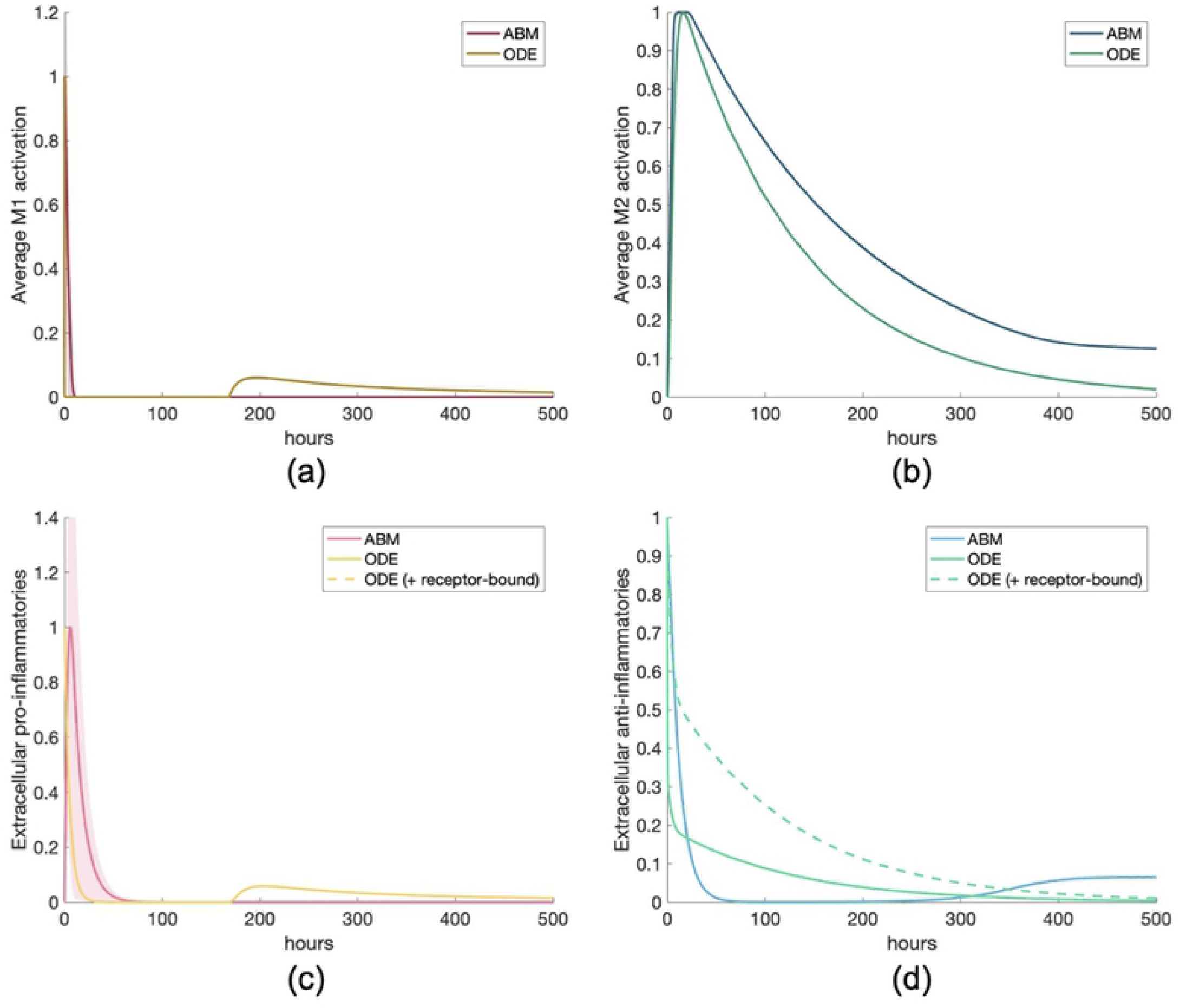
Scenario 1: Simulation of single response to an anti-inflammatory stimulus. All transients are scaled individually by their maximums. (a) M1 activation, (b) M2 activation, (c) PIM, (d) AIM.

Figure 6(c) shows that some PIM is produced in the ABM due to a small percentage of M1 activation existing in the naive macrophages (see Figure 3 to see how naive macrophages are defined), but both models show a decrease to zero in the presence of a large concentration of AIM. Similarly to the calibrating experiment in Figure 5, we show M1 activation in relation to M2 activation in Figure 7 to better visualize the magnitude of the pro-inflammatory response, which is very small in relation to the much larger anti-inflammatory stimulus.

**Fig 7.**
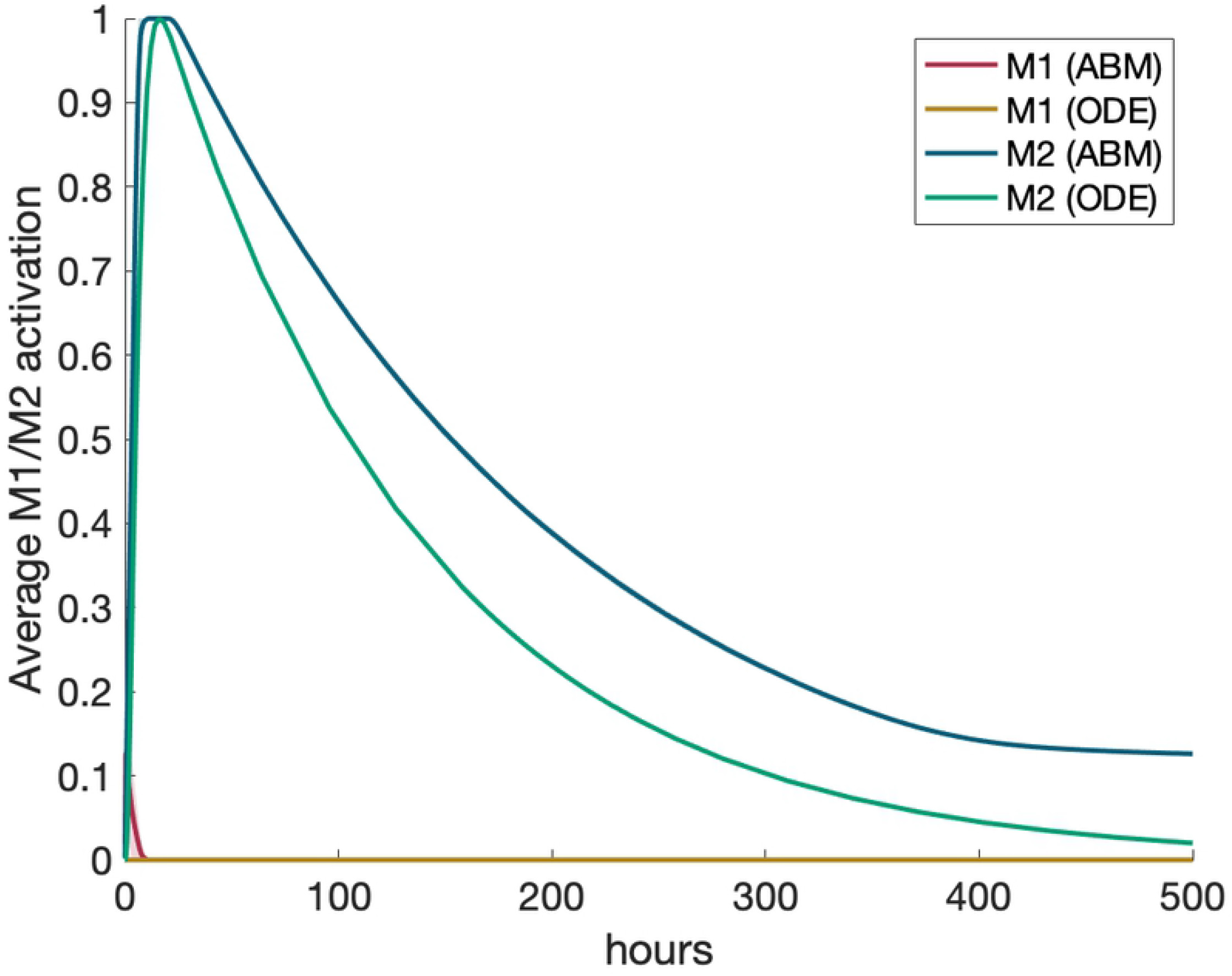
For anti-inflammatory stimulus, M1 and M2 activation are comparable. Scenario 1: M1 and M2 activation response to an anti-inflammatory stimulus. ODE and ABM results are scaled by the maximum M2 activation to compare to maximum M1 activation, which is nearly nonexistent in comparison to M2.

### Scenario 2: Multiple macrophages with pro-inflammatory stimulus

We then introduced recruitment/turnover and cell lifespan. In the ABM, the grid was expanded to 40-by-40 with ten M0 macrophages initially, and the recruitment feature was turned on. Naive and activated macrophages were randomly assigned lifespans of 24 ± 6 and 12 ± 3 hours, respectively. In the ODE, all ten macrophage compartments were utilized and had lifespans of 12 ± 3 hours. In this scenario, we introduced a large pro-inflammatory stimulus into the model. Results are shown in Figure 8.

**Fig 8.**
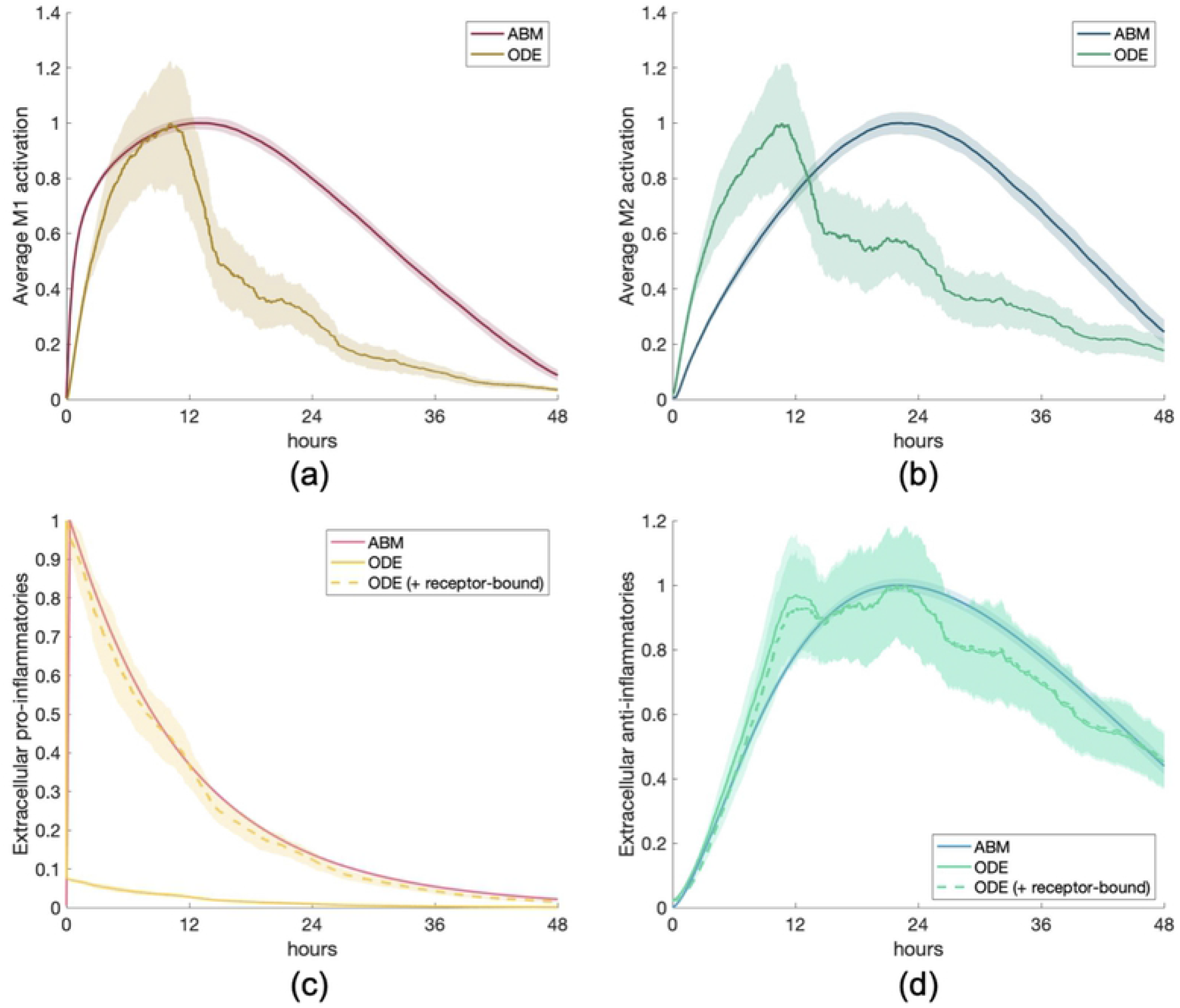
Multi-cell response to a pro-inflammatory stimulus. Scenario 2: M1/M2 response to model of multiple macrophages, activated by an initial amount of pro-inflammatory mediators. All transients are scaled individually by their maximums. (a) M1 activation, (b) M2 activation, (c) PIM, (d) AIM.

In this scenario, M1 and M2 activation in the ODE occur before the ABM, despite similar dynamics for the anti-inflammatory mediators between the two models (panel (d)). Including receptor-bound TNF*α* (Figure 8(c)) makes a significant difference in the dynamics. Our ODE model shows that when naive macrophages are introduced into an environment with a high concentration of TNF*α*, receptors quickly bind to free TNF*α*. Therefore, extracellular TNF*α* in the ODE did not compare well to PIM in the ABM, since receptors are not modeled in the ABM. When receptor-bound TNF*α* was added to the PIM total shown in Figure 8, the dynamics matched up almost perfectly to the ABM. Figure 8(d) shows almost no difference between extracellular IL-10 only and extracellular IL-10 with receptor-bound IL-10, suggesting that accounting for both populations matters more when a large amount of extracellular mediators is introduced rather than the resulting dynamics are observed over time. Standard deviations, shown as the shaded regions in the figures, are also higher than the single-macrophage simulations, since cell lifespan and recruitment provide additional randomness.

### Scenario 3: Multiple macrophages with anti-inflammatory stimulus

The same initial conditions were used for this scenario as in the previous one, except instead of a pro-inflammatory stimulus, an anti-inflammatory stimulus was introduced into the system. Results are shown in Figure 9. PIM and M1 activation were very small compared to AIM and M2 activation, so we do not show their dynamics.

**Fig 9.**
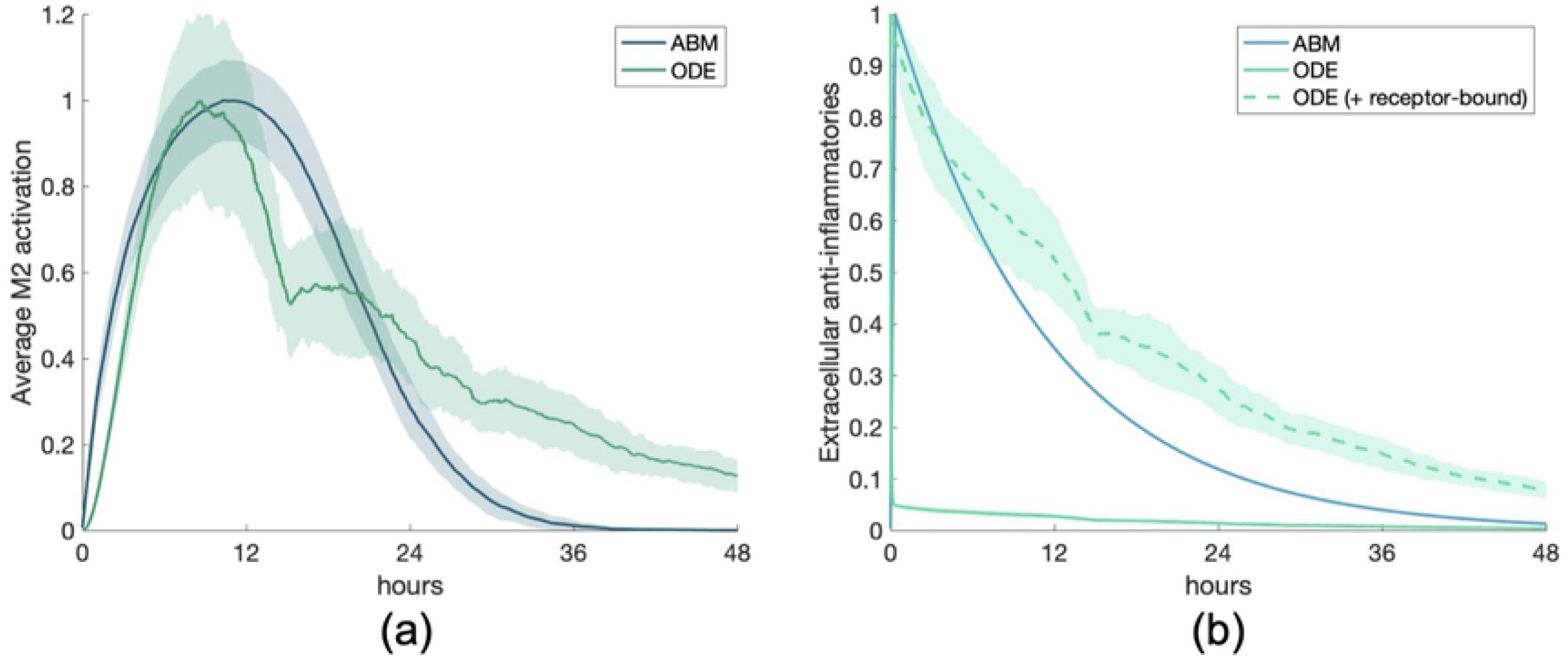
Multi-cell response to anti-inflammatory mediators. Scenario 3: M1/M2 response to M1 macrophages activated by an initial amount of anti-inflammatory mediators. All transients are scaled individually by their maximums. (a) M2 activation, (b) AIM.

M2 activation in the two models are very similar, with the ODE showing a slightly longer tail after the peak of activation. This is paired with a slower decrease of IL-10 (AIM) when receptor-bound IL-10 is taken into account, similarly to PIM in the previous scenario. For a large anti-inflammatory stimulus, similar dynamics are observed between both models when the ODE transient includes receptor-bound IL-10.

### Scenario 4: M1 macrophages with anti-inflammatory stimulus

Next we examined what would happen to an M1 environment when an anti-inflammatory stimulus is introduced into the system. We first needed to determine what this M1 environment would look like as initial conditions that could be used to begin the simulation.

For the ODE, we set all ten macrophages to an M1 phenotype based on the maximum activation that occurs in the calibrating experiment. This maximum occurs around hour 13, so we used the variable values at this time as the initial conditions for all macrophages. We then added a high concentration of IL-10 (the same amount as in Scenario 3) and ran the simulation.

For the ABM, M1 macrophages are defined as having M1_act_ > 0.5 and produce pro- and anti-inflammatory mediators proportional to their activation. To account for recruitment, the equivalent of which in the ODE model is turnover to naive initial conditions, we introduced into the system the number of M1 macrophages at the time when M1 activation was at its highest in Scenario 2. We find that this occurred roughly at hour 12, when there were 205 macrophages. We used this number of M1 macrophages as the initial conditions, along with the same amount of anti-inflammatory mediators as in Scenario 3. We performed two simulations to account for receptor-bound TNF*α* - in the first simulation, we started without any extracellular TNF*α*. Second, we considered no TNF*α* to also include no receptor-bound TNF*α*. Figure 10 shows the results for average activation and extracellular mediators.

**Fig 10.**
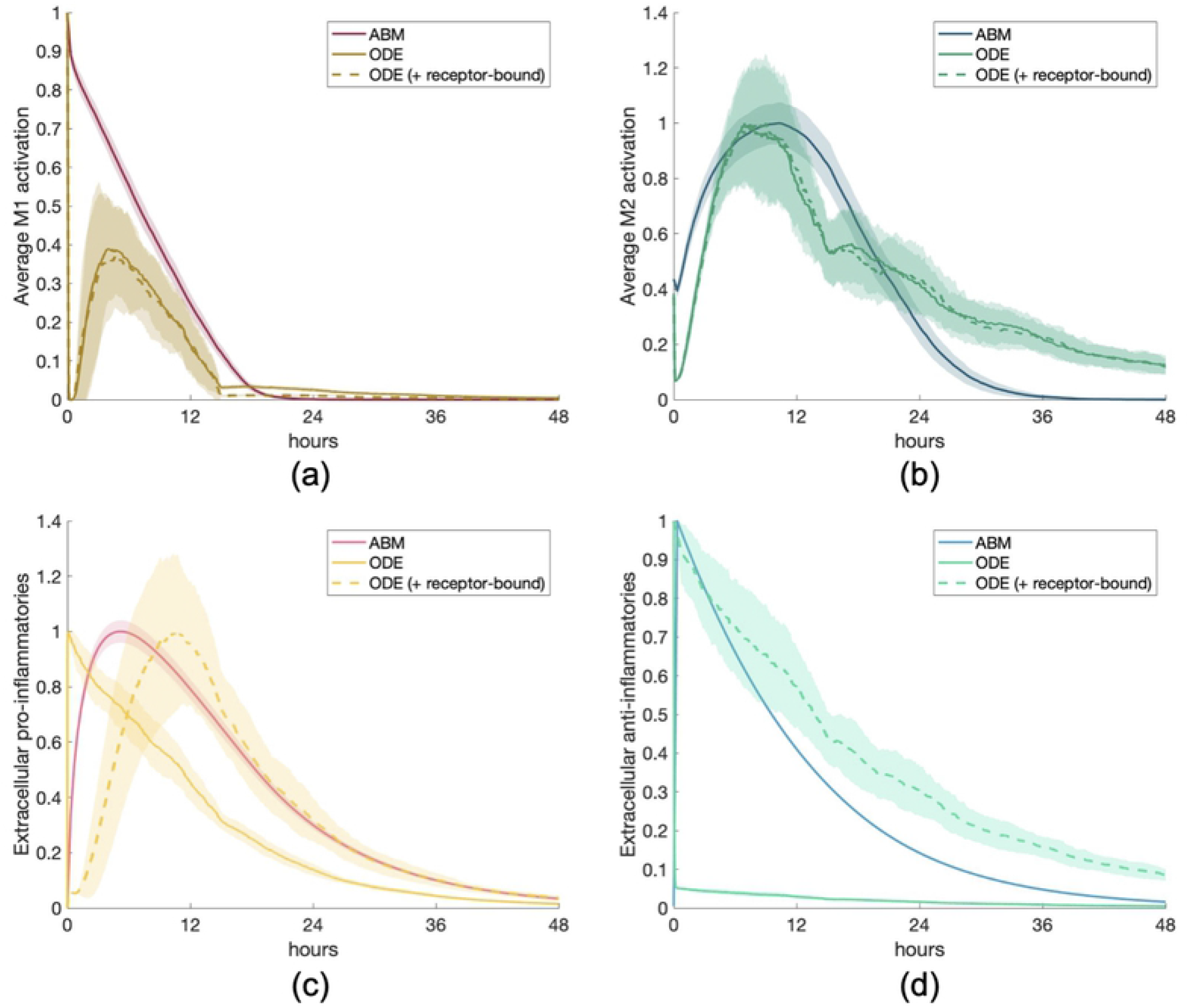
Multi-cell response to AIM in M1 environment. Scenario 4: M1/M2 response to anti-inflammatory stimulus introduced into an M1-polarized system. Transients are scaled individually by their maximums. (a) M1 activation, (b) M2 activation, (c) PIM, (d) AIM.

M2 activation is similar, with the tail of M2 activation and AIM slightly longer in the ODE than the ABM. AIM have a similar response as in the previous scenario, such that with a large anti-inflammatory stimulus, including receptor-bound IL-10 in the AIM improve the ODE model’s similarity to the ABM dynamics. Similarly, including receptor-bound TNF*α* in the total PIM matches ABM dynamics better, though in this case PIM production increases at a slightly higher rate in the ABM. Also, M1 activation shows a small rebound before it decreases to zero. Since AIM do not stimulate the pro-inflammatory signaling pathway but rather inhibit it, this rebound may be due to residual NF*κ*B and TNF*α* in the cytosol and nucleus of the M1 macrophages, taking some time to make its way downstream before being used to produce a small amount of extracellular TNF*α*.

### Scenario 5: Half M1 and half M2

We then observed the results of initializing the models to a state of high activation such that half of the macrophages present were activated to an M1 phenotype and half were M2.

For the ODEs, we used the same initial conditions for M1 macrophages as in Scenario 4, and used a similar method to obtain initial conditions for M2. Hour 16 in Scenario 1 was the time around which peak M2 activation occurs. Five macrophages had M1 initial conditions and the other five had M2 initial conditions.

For the ABM, we used a total of 200 macrophages to represent a state of high activation, similar to the maximum amount of macrophages in Scenario 4. Half were defined as M1 and half as M2. Figure 11 shows the results for M1 and M2 activation and for pro- and anti-inflammatory mediators. When receptor-bound mediators were not taken into account, only extracellular TNF*α* and IL-10 were set to zero at the beginning of the simulation. When receptor-bound mediators were considered part of the overall TNF*α* and IL-10 concentrations, they were also set to zero.

**Fig 11.**
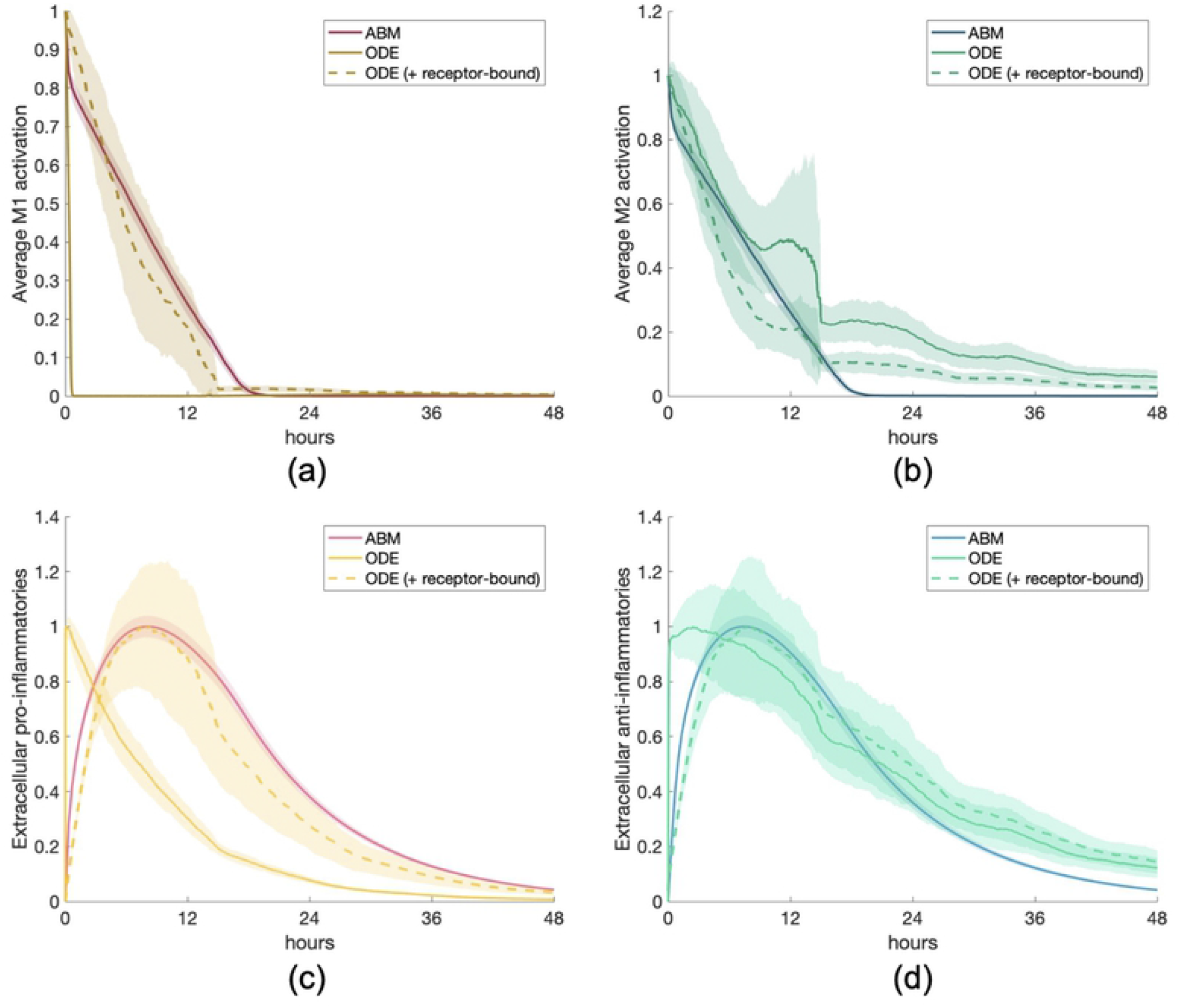
Multi-cell response to mixed M1/M2 environment. Scenario 5: M1/M2 response to a state of activation in which half of the macrophages present are M1 macrophages and half are M2. Transients are scaled individually by their maximums. (a) M1 activation, (b) M2 activation, (c) PIM, (d) AIM.

The ODE model results that included receptor-bound mediators in the total were more similar to ABM results, reflected in all four panels of Figure 11. M1 and M2 activation decay at similar rates due to low production of mediators, and the maximum PIM and AIM show similar behavior as well. The ODE model has consistently shown a longer tail in the overall anti-inflammatory response, both in AIM and M2 activation.

### Scenario 6: PIM activation with wash and anti-inflammatory stimulus

It is common in experimental setups to perform a wash, where cells are treated with a stimulus, then “washed” with a solution to remove external mediators [31]. We replicated this experiment by beginning with the same initial conditions as in Scenario 2: 10 naive macrophages and a pro-inflammatory stimulus. Then at hour 12, the cells, at whatever state they were in at that time, were ”washed” such that PIM and AIM were set to zero and a high amount of AIM was added (same as initial amount in Scenario 3). In the case of considering receptor-bound mediators, receptor-bound TNF*α* was also set to zero at hour 12. Results are shown in Figure 12. Since the times at which M1 and M2 activation is affected most by the wash is different for the ABM versus the ODE model, we compare experiments to examine how they differ from a control, where there is no wash and no AIM added at 12 hours. Therefore, we show four cases, all of which are initialized with PIM: 1) PIM with no later intervention, 2) no wash, AIM added at 12 hours, 3) wash with AIM added, 4) wash with no AIM added. In the future, experiments could be performed with data collected when the models’ dynamics differ significantly in order to select which model best replicates the experimental results.

**Fig 12.**
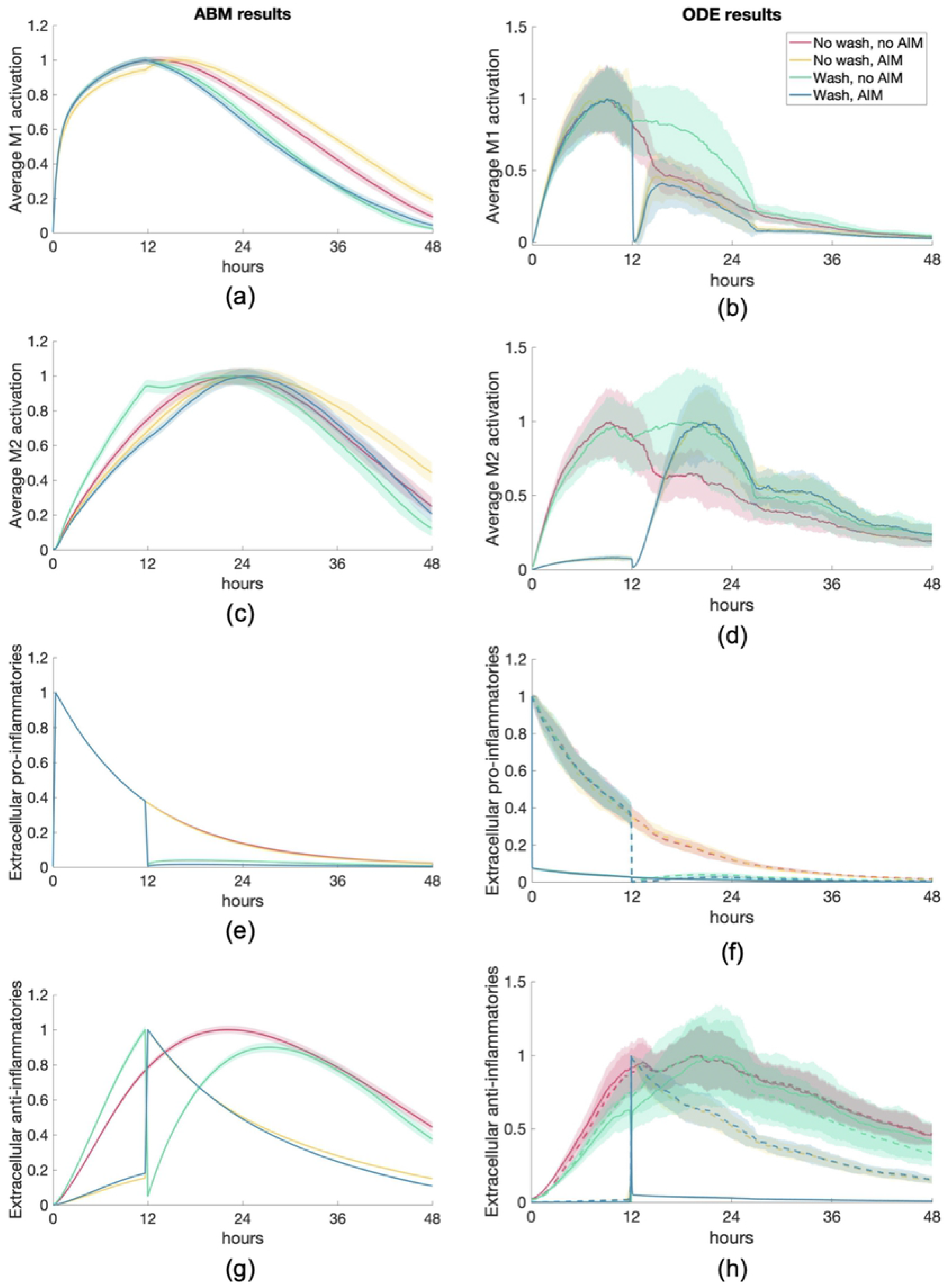
*In silico* simulation of a wash experiment. Scenario 6: M1/M2 response to an initial pro-inflammatory stimulus and either wash or no wash, with AIM added or not added at hour 12. Transients are scaled individually by their maximums. Column 1: ABM results. Column 2: ODE results. (a, b) M1 activation, (c, d) M2 activation, (e, f) PIM, (g, h) AIM.

Figure 12 shows that M1 and M2 activation in the ABM responds similarly regardless of the experiment, whereas they have more distinct results in the ODE simulations. The ODE model has a more immediate response to the AIM than the wash, shown in the sharp changes at hour 12 for the blue and yellow curves in panels (b) and (d). On the other hand, activation in the ABM does not show these sharp changes; rather, they are more gradual even though large jumps are reflected in the PIM and AIM dynamics. Incorporating the receptor-bound mediators into the extracellular AIM and PIM in the ODE simulations shows nearly an exact match with the ABM results in panels (e) through (h). Furthermore, in both model types, M1 activation generally peaks before M2 activation. One noticeable difference is that with the wash, no AIM experiment, more time is needed in the ABM to return to its original levels whereas the ODE model shows a faster rebound. Examining these four scenarios allowed us to observe how the models respond to different variations of stimuli and pinpoint the sensitivity of both models to these stimuli. It could also aid in selecting the best way to create an *in silico* representation of an experiment such as a wash.

### Sensitivity analysis results

Through performing a sensitivity analysis, we were able to determine model parameters to which M1/M2 activation and concentrations of PIM and AIM are most sensitive. We also compared the results between the two models and distinguish whether these key parameters are model-specific or common to both. Results for the single-macrophage ABM are shown in Figure 13. Although we obtained results for 50 and 100 hours, percent change from the original parameter values was very low; therefore, they are not included in Figure 13. Furthermore, we only show results for parameters with a percent change greater than 50% for at least one variable and time point.

**Fig 13.**
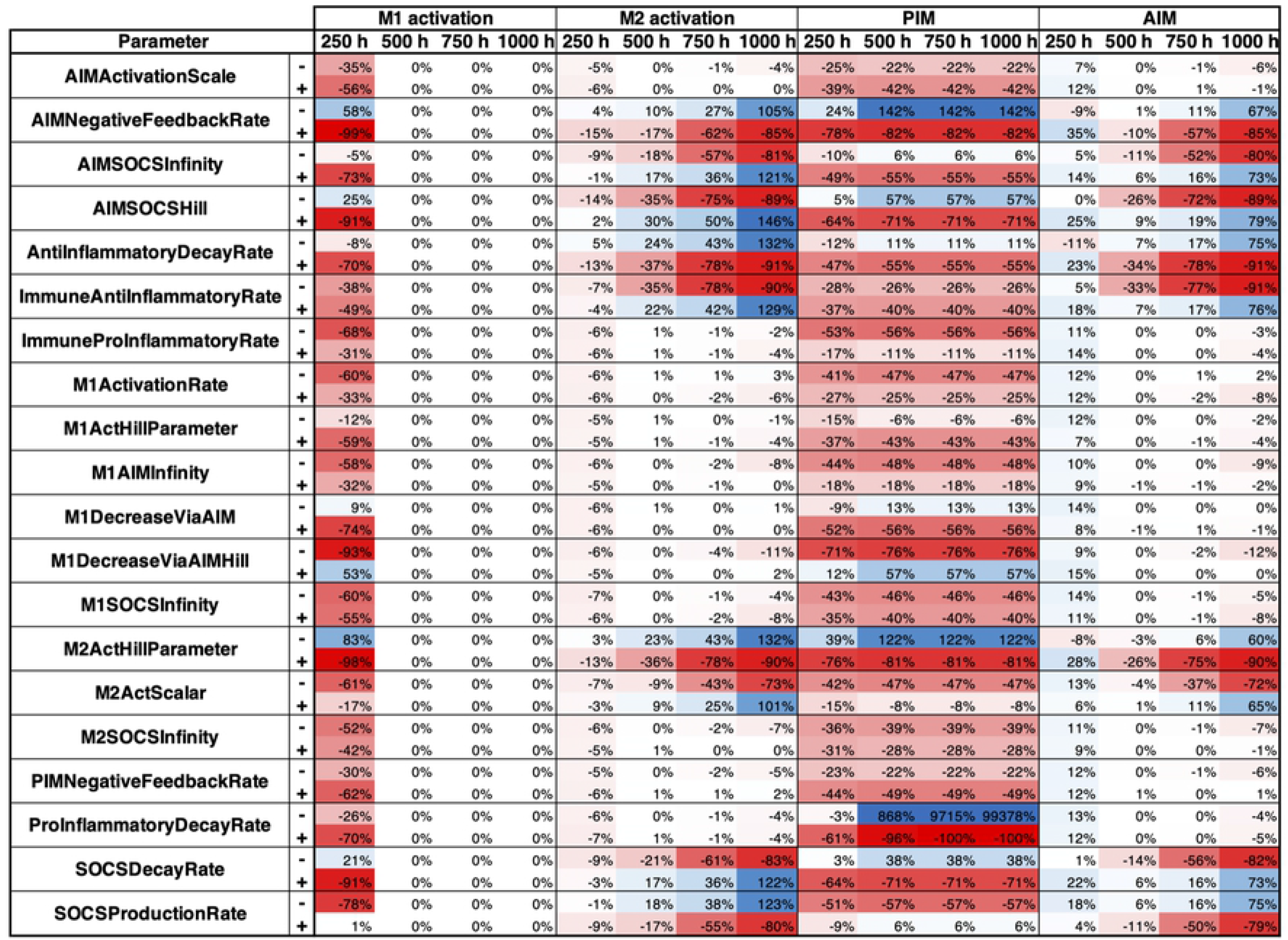
Sensitivity analysis results for single-macrophage ABM. Parameters were perturbed by increasing or decreasing the original value by 10%, shown as “+” and respectively, next to each parameter name. Parameters with 50% change or more for at least one variable and time point are shown.

**Fig 14.**
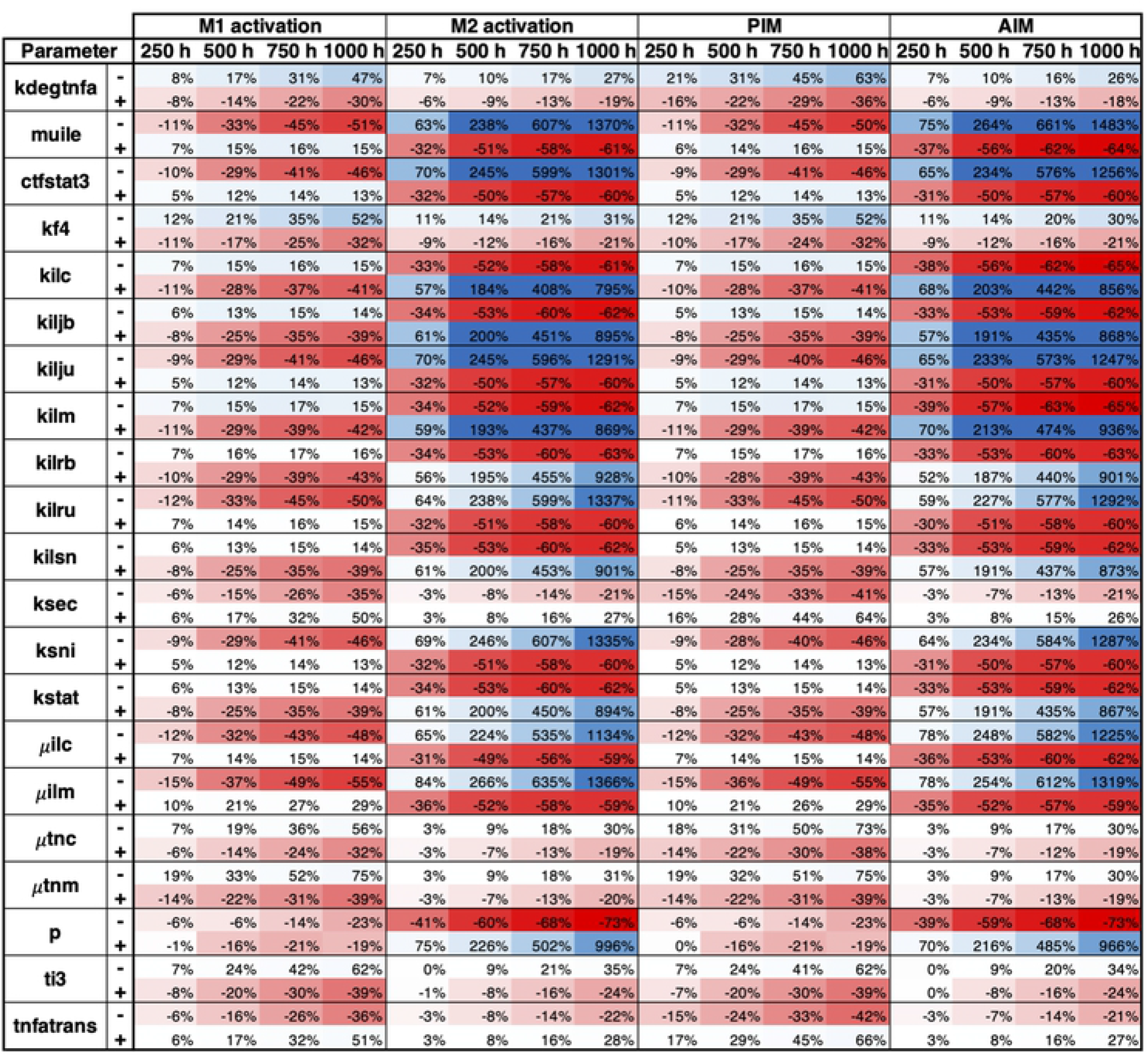
Sensitivity analysis results for single-macrophage ODE model. Parameters were perturbed by increasing or decreasing the original value by 10%, shown as “+” and “-”, respectively, next to each parameter name. Parameters with 50% change or more for at least one variable and time point are shown.

Of the 56 parameters in the ODE model, 21 show changes by more than 50% for at least one time point and variable tested. M2 activation and AIM changed the most when parameters were perturbed, particularly at 500, 750, and 1000 hours. The majority of these sensitive parameters are related to extracellular or intracellular IL-10, relating to extracellular IL-10 binding and unbinding to and from its receptor and the step-by-step creation of the IL-10-STAT3 complex with JAK1 and Tyk2. A number of other highly sensitive parameters are found in TNF*α*-related equations, relating to its activation and transcription, such as *μ_tnc_, μ_tnm_, k_sec_*, and *k_tnfatrans_*. Not many parameter relating to the signaling pathways involving I*κ*B*α* and NF*κ*B were sensitive, pointing to the possibility that these steps do not need to be modeled explicitly.

In the ABM, 20 parameters result in changes by more than 50% for at least one variable and one time point. M1 activation is very sensitive at the first time point tested, 250 hours, and not sensitive at all for the later time points. This is due to the shorter time course of M1 activation in the ABM, where M1 activation returns to 0 within about 250 hours and does not increase again, in contrast to the ODE model where M1 activation decreases more gradually after the peak. Furthermore, the 0% sensitivity after 250 hours in the ABM highlights the discrete nature of species in this type of model; once M1 macrophages disappear from the grid, they do not return. In contrast, PIM, a continuous variable that diffuses throughout the grid, shows high sensitivity at all time points measured, even though PIM appears to be nearly zero in Figure 4(c). This highlights how the type of model can affect sensitivity results. Furthermore, in contrast to the ODE model, the highly sensitive parameters found from the ABM do not focus on any particular mechanisms; rather, they reflect a variety of steps in the immune response, from M1 and M2 activation (M1ActivationRate, M2ActHillParameter, M2ActScalar) to SOCS activation and regulation SOCS (AIMSOCSInfinity, M2SOCSInfinity, SOCSProductionRate).

## Discussion

With still much unknown about M1-M2 polarization and the important role it plays in the pathogenesis of many diseases [5], our modeling approaches and scenarios contribute to the body of knowledge surrounding macrophage polarization by providing a comparison of *in silico* platforms to test hypotheses and highlight mechanisms that may be necessary or unnecessary to include in future models.

By using the same basic principles of M1/M2 activation, interaction with mediators, and cell lifespan, our two distinctly different models provided surprisingly similar results after tuning to a common calibrating experiment. In particular, peak times and overall shapes of the transients were similar in most cases. Whereas our ODE model accounted for relatively detailed subcellular signaling, where each term represented a different interaction within the cell as well as with extracellular mediators, our ABM simplified the interactions to reflect similar roles of M1/M2 activation without the detail of individual mechanisms and interactions. Rather, only M1/M2 activation and mediators were measured in the model.

A common difference between models was a longer tail of M2 activation and AIM activity across several scenarios. This was also seen in the calibrating scenario, where M2 activation decreases more quickly in the ABM. Future work could include finer tuning of the parameters to better align the model results.

Another thread throughout this work is the consideration of receptor-bound TNF*α* and IL-10 in the ODE model. In most scenarios, especially those with high amounts of one mediator (Scenarios 2-4) or both (Scenarios 5-6), incorporating receptor-bound mediators into the overall concentration of mediators improved similarity to the ABM results. Though this disparity was initially unexpected, it was not surprising since the ABM does not explicitly model receptors such that extracellular mediators are not removed from the population when they interact with a macrophage. Due to the significant difference when taking into account receptors versus not taking them into account, future changes to the ABM may involve accounting for receptor-bound mediators by explicitly including receptors or PIM and AIM in the extracellular population could be decreased when they come into contact with a macrophage, representing binding to receptors.

We also wanted to examine the differences that incorporating space (ABM) or detailed subcellular signaling (ODEs) would make in the resulting dynamics. A notable difference between the two models is seen in Figure 10(a), where residual amounts of M1-related variables such as intracellular forms of NF*κ*B and TNF*α* resulted in a small downstream bump in M1 activation in the ODE, whereas the ABM, which does not account for these variables, showed a more gradual, constant decrease of M1 activation to zero. Another significant difference was observed in the “wash” experience in Scenario 6, where the ODE model had a greater sensitivity to immediate changes in PIM and AIM than the ABM. In the ABM, rules of macrophage activation are defined such that activation decreases gradually when a stimulus is not present, whereas in the ODE model the explicit transcription of mRNA responds directly and more immediately to a lack of extracellular mediators. Interestingly, this discrepancy did not significantly affect the other scenarios. This is an area of future investigation, especially if these models could be validated with experimental data. Overall, the incorporation of multi-step subcellular signaling was not very important since the ABM did not include subcellular signaling and we obtained very similar dynamics from both models.

We did not observe significant differences regarding the spatial dynamics of the ABM versus the well-mixed assumption of the ODE model, although this was not a focus of our analysis. It has been shown in previous ABMs involving macrophages, such as modeling granuloma formation in tuberculosis [32], that incorporating the ability of macrophages to interact on a spatial level and gather together is important to the immune response. Future simulations and scenarios could involve putting initial amounts of PIM and AIM on different areas within the grid or in different patterns to observe more carefully how space plays a role in M1/M2 activation.

Our sensitivity analysis showed that in the ODE model, M1 & M2 activation, PIM, and AIM were not highly sensitivity to parameters relating to the NF-κB pathway. This suggests that these mechanisms may not need to be modeled explicitly. In the ABM, the mechanisms reflected in the highly sensitive parameters were more evenly distributed among each step of the series of rules, suggesting that the mechanisms included were each necessary to capture the dynamics of the variables of interest. Additional understanding of uncertainty in the two models could be ascertained by examining how sensitivity differs between scenarios, and how the discrete aspects of the ABM contribute to the sensitivity profiles of the variables of interest.

Based on our findings from comparing the two models, we recommend a focus on the main interactions of extracellular mediators and macrophages, where M1/M2 polarization can occur on a continuous spectrum, reflecting the current knowledge and modeling practices of macrophage activation [9, 33, 34]. Important feedback loops in the pro- and anti-inflammatory phases of the immune response are: the positive feedback loop of M1 activation, upregulation of M2 via M1, and the negative feedback loop in which M2 decreases both M1 and itself. Initially, our ABM did not include SOCS, a family of intracellular proteins produced by the IL-10 pathway to regulate itself. Without this regulatory feedback loop, M2 activation and AIM did not decrease back to its initial state, but when we added SOCS to the ABM, we obtained the expected dynamics such that the calibrating experiment results of the ABM were similar to the ODE model, which did include SOCS. Whether these interactions and feedback loops are modeled explicitly through signaling pathways or through general rules was less important for our purposes, as our results from the two approaches were similar, as long as they were included in some manner.

Future work necessary to confirm our hypotheses via the scenarios described above is to fit both models, especially the calibrating experiment, to additional data. Thus far, the ODE model based on LPS-induced dynamics utilized data to fit the parameters. More sophisticated parameter estimation methods, such as obtaining correlations between parameters and a sensitivity analysis, would be useful due to the large number of parameters in the model. Furthermore, currently both models are meant to represent the immune response to a general insult. These models can be adapted to incorporate the key players and mechanisms involved in specific injuries such as bacterial or viral infections, wound healing, sepsis, or COPD.

## Acknowledgments

This work was supported by the National Institutes of Health via award R21HL146250 (R.H.).

## References

1. Misharin AV, Morales-Nebreda L, Mutlu GM, Budinger GRS, Perlman H. Flow Cytometric Analysis of Macrophages and Dendritic Cell Subsets in the Mouse Lung. American Journal of Respiratory Cell and Molecular Biology. 2013;49(4):503–510. doi:10.1165/rcmb.2013-0086MA.

2. Mahbub S, Deburghgraeve CR, Kovacs EJ. Advanced Age Impairs Macrophage Polarization. Journal of Interferon & Cytokine Research. 2012;32(1):18–26. doi:10.1089/jir.2011.0058.

3. Smith TD, Tse MJ, Read EL, Liu WF. Regulation of macrophage polarization and plasticity by complex activation signals. Integrative Biology. 2016;8(9):946–955. doi:10.1039/c6ib00105j.

4. Gibon E, Loi F, Córdova LA, Pajarinen J, Lin T, Lu L, et al. Aging Affects Bone Marrow Macrophage Polarization: Relevance to Bone Healing. Regenerative Engineering and Translational Medicine. 2016;2(2):98–104. doi:10.1007/s40883-016-0016-5.

5. Bosco MC. Macrophage polarization: Reaching across the aisle? Journal of Allergy and Clinical Immunology. 2019;143(4):1348–1350. doi:10.1016/j.jaci.2018.12.995.

6. Maiti S, Dai W, Alaniz RC, Hahn J, Jayaraman A. Mathematical Modeling of Pro- and Anti-Inflammatory Signaling in Macrophages. Processes. 2014;3(1):1–18.

7. Moya C, Huang Z, Cheng P, Jayaraman A, Hahn J. Investigation of IL-6 and IL-10 signalling via mathematical modelling. IET Systems Biology. 2011;5(1):15–26. doi:10.1049/iet-syb.2009.0060.

8. Frank AS, Larripa K, Ryu H, Snodgrass RG, Röblitz S. Bifurcation and sensitivity analysis reveal key drivers of multistability in a model of macrophage polarization. Journal of Theoretical Biology. 2021;509:110511.

9. Zhao C, Mirando AC, Sove RJ, Medeiros TX, Annex BH, Popel AS. A mechanistic integrative computational model of macrophage polarization: Implications in human pathophysiology. PLOS Computational Biology. 2019;15(11):e1007468. doi:10.1371/journal.pcbi.1007468.

10. Rex J, Albrecht U, Ehlting C, Thomas M, Zanger UM, Sawodny O, et al. Model-Based Characterization of Inflammatory Gene Expression Patterns of Activated Macrophages. PLOS Computational Biology. 2016;12(7):e1005018. doi:10.1371/journal.pcbi.1005018.

11. Kirschner D, Pienaar E, Marino S, Linderman JJ. A review of computational and mathematical modeling contributions to our understanding of Mycobacterium tuberculosis within-host infection and treatment. Current Opinion in Systems Biology. 2017;3:170–185. doi:10.1016/j.coisb.2017.05.014.

12. Nickaeen N, Ghaisari J, Heiner M, Moein S, Gheisari Y. Agent-based modeling and bifurcation analysis reveal mechanisms of macrophage polarization and phenotype pattern distribution. Scientific Reports. 2019;9(1):12764. doi:10.1038/s41598-019-48865-z.

13. Reynolds A, Rubin J, Clermont G, Day J, Vodovotz Y, Ermentrout GB. A reduced mathematical model of the acute inflammatory response: I. Derivation of model and analysis of anti-inflammation. Journal of theoretical biology. 2006;242(1):220–236.

14. Torres M, Wang J, Yannie PJ, Ghosh S, Segal RA, Reynolds AM. Identifying important parameters in the inflammatory process with a mathematical model of immune cell influx and macrophage polarization. PLoS computational biology. 2019;15(7):e1007172.

15. Cell Signaling | Learn Science at Scitable;. Available from: https://www.nature.com/scitable/topicpage/cell-signaling-14047077/.

16. Translation: DNA to mRNA to Protein | Learn Science at Scitable;. Available from: https://www.nature.com/scitable/topicpage/translation-dna-to-mrna-to-protein-393/.

17. Wang N, Liang H, Zen K. Molecular Mechanisms That Influence the Macrophage M1–M2 Polarization Balance. Frontiers in Immunology. 2014;5. doi:10.3389/fimmu.2014.00614.

18. Carey AJ, Tan CK, Ulett GC. Infection-induced IL-10 and JAK-STAT. JAK-STAT. 2012;1(3):159–167. doi:10.4161/jkst.19918.

19. Lipniacki T, Paszek P, Brasier AR, Luxon B, Kimmel M. Mathematical model of NF-κB regulatory module. Journal of Theoretical Biology. 2004;228(2):195–215. doi:10.1016/j.jtbi.2004.01.001.

20. Dimitriou ID, Clemenza L, Scotter AJ, Chen G, Guerra FM, Rottapel R. Putting out the fire: coordinated suppression of the innate and adaptive immune systems by SOCS1 and SOCS3 proteins. Immunological Reviews. 2008;224(1):265–283. doi:10.1111/j.1600-065X.2008.00659.x.

21. Qasimi P, Ming-Lum A, Ghanipour A, Ong CJ, Cox ME, Ihle J, et al. Divergent Mechanisms Utilized by SOCS3 to Mediate Interleukin-10 Inhibition of Tumor Necrosis Factor α and Nitric Oxide Production by Macrophages. Journal of Biological Chemistry. 2006;281(10):6316–6324. doi:10.1074/jbc.M508608200.

22. Rawlings JS, Rosler KM, Harrison DA. The JAK/STAT signaling pathway. Journal of Cell Science. 2004;117(8):1281–1283. doi:10.1242/jcs.00963.

23. Sabat R, Grutz G, Warszawska K, Kirsch S, Witte E, Wolk K, et al. Biology of interleukin-10. Cytokine & Growth Factor Reviews. 2010;21(5):331–344. doi:10.1016/j.cytogfr.2010.09.002.

24. Riley JK, Takeda K, Akira S, Schreiber RD. Interleukin-10 Receptor Signaling through the JAK-STAT Pathway: Requirement for Two Distinct Receptor-Derived Signals for Anti-Inflammatory Action. Journal of Biological Chemistry. 1999;274(23):16513–16521. doi:10.1074/jbc.274.23.16513.

25. Croker BA, Kiu H, Nicholson SE. SOCS regulation of the JAK/STAT signalling pathway. Seminars in Cell & Developmental Biology. 2008;19(4):414–422. doi:10.1016/j.semcdb.2008.07.010.

26. Tamiya T, Kashiwagi I, Takahashi R, Yasukawa H, Yoshimura A. Suppressors of Cytokine Signaling (SOCS) Proteins and JAK/STAT Pathways. Arteriosclerosis, Thrombosis, and Vascular Biology. 2011;31(5):980–985. doi:10.1161/ATVBAHA.110.207464.

27. Yasukawa H, Misawa H, Sakamoto H, Masuhara M, Sasaki A, Wakioka T, et al. The JAK binding protein JAB inhibits Janus tyrosine kinase activity through binding in the activation loop. The EMBO Journal. 1999;18(5):1309–1320. doi:10.1093/emboj/18.5.1309.

28. Yagil Z, Nechushtan H, Kay G, Yang CM, Kemeny DM, Razin E. The enigma of the role of Protein inhibitor of Activated STAT3 (PIAS3) in the immune response. Trends in Immunology. 2010;31(5):199–204. doi:10.1016/j.it.2010.01.005.

29. Hutchins AP, Diez D, Miranda-Saavedra D. The IL-10/STAT3-mediated anti-inflammatory response: recent developments and future challenges. Briefings in Functional Genomics. 2013;12(6):489–498. doi:10.1093/bfgp/elt028.

30. Koga T, Kuwahara I, Lillehoj EP, Lu W, Miyata T, Isohama Y, et al. TNF-α induces MUC1 gene transcription in lung epithelial cells: its signaling pathway and biological implication. American Journal of Physiology-Lung Cellular and Molecular Physiology. 2007;293(3):L693–L701. doi:10.1152/ajplung.00491.2006.

31. Wang Y, Wang YP, Zheng G, Lee VWS, Ouyang L, Chang DHH, et al. Ex vivo programmed macrophages ameliorate experimental chronic inflammatory renal disease. Kidney International. 2007;72(3):290–299. doi:10.1038/sj.ki.5002275.

32. Marino S, Cilfone NA, Mattila JT, Linderman JJ, Flynn JL, Kirschner DE. Macrophage Polarization Drives Granuloma Outcome during Mycobacterium tuberculosis Infection. Infection and Immunity. 2015;83(1):324–338. doi:10.1128/IAI.02494-14.

33. Mosser DM, Edwards JP. Exploring the Full Spectrum of Macrophage Activation. Nature Reviews Immunology. 2008;8(12):958–969. doi:10.1038/nri2448.

34. Marku M, Verstraete N, Raynal F, Madrid-Mencia M, Domagala M, Fournie JJ, et al. Insights on TAM Formation from a Boolean Model of Macrophage Polarization Based on In Vitro Studies. Cancers. 2020;12(12):3664. doi:10.3390/cancers12123664.

